# Tissue Context Drives Regional Differences in Trophoblast Differentiation in Fetal Membranes and Chorionic Villi

**DOI:** 10.1101/2024.11.19.624345

**Authors:** Liheng (Henry) Yang, Allyson Caldwell, Carolyn B. Coyne

## Abstract

Trophoblasts within the fetal membrane primarily serve as protective barriers, shielding the developing fetus from the external environment, while those in chorionic villi facilitate interactions between the maternal and fetal compartments. Although these distinct trophoblast populations perform specialized roles based on their placental location, the mechanisms governing their development and differentiation remain largely unknown. In this study, we derived trophoblast organoids from both the smooth chorion of fetal membranes and chorionic villi from matched human placentas to create parallel *in vitro* models of these distinct trophoblast populations. Using comparative transcriptional profiling with both bulk and single-cell RNA sequencing, we identified subtle transcriptional variations, while overall gene expression patterns and cellular composition remained highly conserved. Comparative single-cell RNA sequencing and differentiation trajectory analysis of organoids derived from chorion and villous tissues, along with their respective tissue counterparts, revealed region-specific gene expression patterns that were only partially conserved in organoids. The similarity between these *in vitro* models suggests that regional differences in trophoblast expression observed *in vivo* are driven by environmental and regional cues. These findings highlight the critical role of local environmental factors in shaping trophoblast function and offer insights into the conserved mechanisms that support placental integrity and fetal development, while emphasizing the influence of region-specific cues *in vivo*.

## Introduction

The placenta forms very early in pregnancy. By the end of the first trimester, the chorionic sac differentiates to form the chorion frondosum (villi-containing portion of the placenta) and the chorion laeve (smooth chorion), which forms the extra-amniotic membranes. Chorionic villi are covered by the syncytiotrophoblast (STB), a multinucleated trophoblast layer that forms from the fusion of mononuclear cytotrophoblasts. At the terminal ends of villous columns, extravillous trophoblasts (EVTs) invade into the maternal-derived decidua basalis and anchor the chorion frondosum to the uterine lining. Unlike EVTs located on the tips of anchoring villi, chorion-derived trophoblasts do not directly invade the decidua parietalis but do express EVT markers including HLA-G^1^. Comparative single-cell RNA-sequencing (scRNA-seq) studies of chorionic villi and smooth chorion isolated from matched mid-gestation human placentas suggests that there are regional differences in the transcriptomics of trophoblasts comprising these distinct sites, which might impact their function and/or immune regulation^2,3^. For example, smooth chorion-derived trophoblasts (schCTBs) express factors that inhibit EVT invasion^2^, suggesting that the lack of decidual invasion observed *in vivo* is at least partially influenced by differences in the expression of key transcripts between these specialized trophoblast populations.

The mechanisms by which villi- and smooth chorion-derived trophoblasts differentiate to maintain distinct cell populations during pregnancy remain poorly defined. Transcriptional profiling of these sites from mid-gestation tissue suggest that similar progenitor/stem populations exist but may display distinct signatures and/or differentiation pathways^2^. Progenitor cells in chorionic villi have been isolated from the first trimester and recapitulate many properties of trophoblasts isolated from the chorion frondosum^4^. Similarly, organoids developed from these progenitor cells in either the first trimester^5^ or from mid-to-late gestation tissue^6^ also recapitulate the properties of villous-derived trophoblasts. In contrast, the progenitor cell population that replenishes the smooth chorion is less defined. Previous work successfully isolated progenitor cell lines from first trimester chorion tissue that recapitulated properties of schCTBs^7,8^. However, stem cell-derived organoid models of schCTBs have not been described, which may better model the spatial morphology of the differentiated chorion and biologically relevant cell-cell and cell-matrix interactions.

Here, we isolated stem/progenitor cells from the chorion laeve of full-term fetal membrane tissue. In parallel, we developed trophoblast organoids from chorionic villous tissue from matched placentas to determine whether there were differences between trophoblast organoids derived from distinct regions of the placenta. Transcriptional profiling by bulk RNASeq revealed a high degree of transcript overlap between organoids isolated from the chorion frondosum and chorion leave, including the expression of canonical markers of distinct trophoblast populations. To further characterize the cell populations within these organoids, we performed comparative single-cell scRNA-seq. This comparative analysis revealed minimal region-specific gene expression differences. We then integrated the organoid datasets with existing scRNA-seq data from mid-gestation placental tissue, uncovering distinct differentiation trajectories for schCTBs *in vivo*, which were only partially retained in organoids. By establishing matched organoid models from distinct placental regions, this study offers a platform to dissect region-specific trophoblast functions and to further understand their contributions to placental development and related pathologies.

## Results

### Long-term expansion of three-dimensional trophoblast organoids derived from full-term human smooth chorion

To establish matched trophoblast organoids from distinct regions of the placenta, we dissected chorionic villi and smooth chorion from matched full-term human placentas (38–41-week gestation) **(Figure 1A, Figure S1A**). We found that the smooth chorion contained high levels of proliferative cells as assessed by Ki67 immunoreactivity (**Figure S1B**). Because proliferating cells can be associated with stem/progenitor cells, we reasoned that the smooth chorion may contain higher numbers of stem/progenitor cells than those observed previously in full-term chorionic villi^6^. To isolate chorion-enriched stem/progenitor cells, we dissected full-term fetal membranes into chorion and amnion layers and carefully removed any decidua tissue. We homogenized the smooth chorion using similar protocols that we developed previously for the isolation of chorionic villi-derived organoids from full-term tissue^6^. In parallel, we homogenized the amnion using established procedures to isolate amnion epithelial cells (AECs) and confirmed their purity (>90%) by immunostaining for the epithelial cell marker cytokeratin-19 (**Figure S1C**). To establish smooth chorion-derived organoids, stem/progenitor cells isolated from smooth chorion were embedded in Matrigel “domes” and submerged in a cocktail of growth factor-containing media used to grow villi-derived TOs^6^. We observed the rapid growth of chorion organoids (COs) within 2-5 days post-isolation, with hCG detectable in the medium as early as 3-4 days post-isolation (**Figure 1C, top**) and large organoid structures visible by 5-7 days post-isolation (**Figure 1B**). This growth was significantly faster than that observed for trophoblast organoids (TOs) developed from chorionic villous tissue of matched placentas, which required 1-2 weeks for derivation following isolation. By ∼5-10 days post-isolation, large COs were visible as three-dimensional structures (**Figure 1C, bottom**). Following their initial isolation and derivation, COs could be expanded at a 1:4∼6 passaging ratio every 5-7 days, exhibiting similar morphology and growth patterns as TOs. Established CO lines could be expanded long term (>20 passages) without significant changes in growth kinetics, morphology, or other general behavior, cryopreserved, and revived successfully upon thawing (**Figure S1D**). We have successfully developed COs from fifteen full-term placentas with a 100% success rate of derivation. Established CO lines were derived from both male and female tissue (8 male and 7 female).

**Figure 1:**
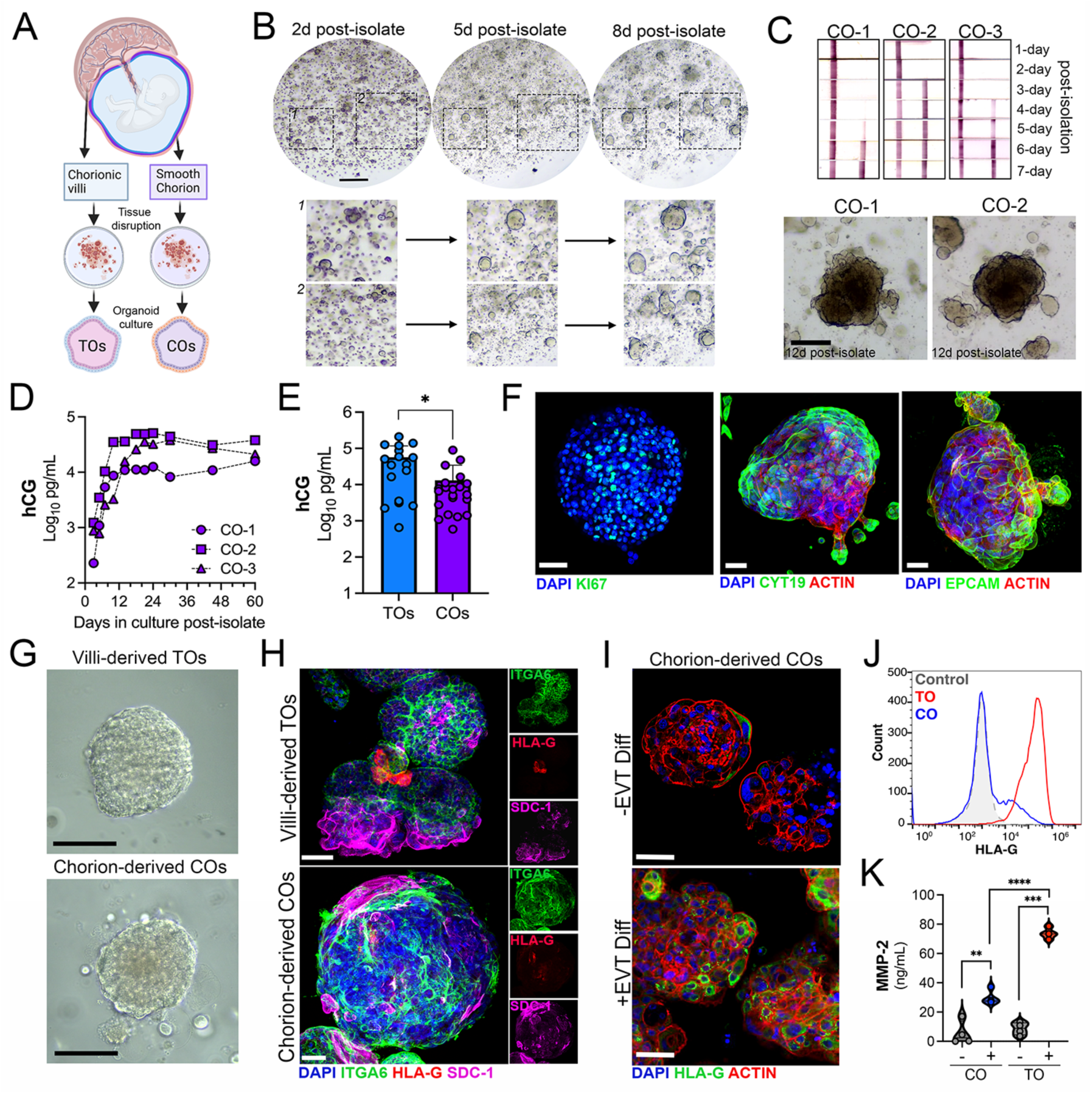
Establishment and characterization of smooth chorion-derived organoids. **(A),** Schematic of the human placenta denoting chorionic villi and smooth chorion used to generate matched organoids. Created using Biorender. **(B)**, Brightfield images of one line of chorion-derived organoids (COs) at 2 days (left), 5 days (middle), and 8 days (right) post-isolation. At bottom, zoomed images of two areas denoted by hatched black boxes at top. Scale bar, 250μm **(C),** Top, representative images of over the counter (OTC) hCG test strips at 1-7 days post-isolation of three independent chorion-derived organoid lines (CO-1, CO-2, and CO-3). Bottom, brightfield images of representative organoids from two lines of chorion-derived organoids (CO-1 and CO-2) at 10 days post-isolation. Scale bar, 100μm. **(D),** hCG levels (shown as log_10_ pg/mL) as determined by Luminex from media collected from three independent CO lines at the indicated days post-isolation. **(E),** hCG levels (shown as log_10_ pg/mL) as determined by Luminex from media collected from matched lines of villous-derived trophoblast organoids (TOs, blue) or matched COs (purple). Each symbol represents an independent well from at least three independent lines. Significance was determined using a student’s t-test (* p<0.05). **(F),** Confocal micrographs of COs immunostained for Ki67 (green, left), cytokeratin-19 (green, middle) and Epcam (green, right). In middle and right, actin is shown in red. In all, DAPI-stained nuclei are in blue. **(G),** Brightfield images of TOs and COs isolated from the same placental tissue. Scale bar, 75μm. **(H),** Confocal micrographs of COs and TOs immunostained for ITGA6 (green), HLA-G (red), and SDC-1 (purple). DAPI-stained nuclei are in blue. Individual channels are shown at right. **(I),** Confocal micrographs of COs cultured in standard (-EVT diff, top) conditions or conditions optimized for EVT differentiation (+EVT diff, bottom) and immunostained for HLA-G (in green). Organoids were counterstained for actin (in red). DAPI-stained nuclei are in blue. **(J),** Flow cytometry analysis of cell surface HLA-G expression in COs (blue) and TOs (red) following EVT differentiation. Isotype control (grey) was included to account for non-specific binding. The x-axis represents fluorescence intensity of HLA-G staining, and the y-axis shows cell count. Data are representative of three independent experiments. **(K),** Levels of MMP-2 (shown as ng/mL) in the supernatant of COs (left) or TOs (right) cultured under standard conditions (-) or conditions optimized for EVT differentiation (+). Each symbol represents an independent sample from at least four replicates. Significance was determined using a student’s t-test (** p<0.01, *** p<0.001, **** p<0.0001). Scale in (F, H, I), 50μm.

### Chorion organoids recapitulate the signature of schCTBs

Because the chorion is composed of schCTBs, we determined whether COs recapitulated the phenotypes of these cells. We confirmed that COs released high levels of hCG, which could be maintained for extended times post-derivation and after repeated passaging (**Figure 1D**). Levels of hCG in media collected from COs was comparable to the levels in media from villi-derived TOs from matched placental tissues, although the levels in TO media were consistently higher than those in COs (**Figure 1E**). Established COs contained high numbers of Ki67^+^ cells as assessed by immunostaining, consistent with their capacity for long-term propagation (**Figure 1F, left**) and also expressed the trophoblast markers cytokeratin-19, and Epcam (**Figure 1F, middle and right**). Established lines of TOs and COs showed similar morphology at comparable passage numbers and culture days post-passage (**Figure 1G**). Confocal microscopy of immunostaining for markers including the STB marker SDC-1, the cytotrophoblast (CTB) marker ITGA6, and the EVT marker HLA-G confirmed that like TOs, COs differentiated to contain distinct trophoblast cell types (**Figure 1H**). To determine if we could enrich for the presence of EVTs in COs, we utilized a previously described two-step EVT differentiation procedure that leads to the significant enrichment of EVTs in TOs^4,6,9^. This procedure led to a significant increase in HLA-G-positive cells in COs (**Figure 1I).** However, EVT differentiation in COs was less pronounced compared to TOs, as assessed by flow cytometry for cell surface HLA-G expression (**Figure 1J**). Consistent with this, we found that EVT differentiation caused an increase in MMP-2 levels, which is specifically secreted by EVTs, in the culture media of TOs compared to that of COs (**Figure 1K**). Collectively, these data demonstrate that established COs recapitulate the hormone release and expression of schCTB markers, are morphologically like TOs, and differentiate into distinct trophoblast lineages.

### Transcriptional profiling of organoids and primary cells derived from distinct placental regions

Since COs closely resembled TOs morphologically and expressed markers of distinct trophoblast cell types, we next performed bulk RNA-seq on established CO and TO lines to compare their transcriptional signatures. To ensure the purity of COs and rule out contamination by decidua-derived cells, we also conducted RNA-seq profiling of decidua organoids (DOs) derived from matched placental tissue. Finally, we compared the transcriptional profiles of COs with amnion epithelial cells (AECs) to identify differences in gene expression between cells comprising distinct layers of the fetal membrane and further validate the purity of the CO preparations. We performed comparative expression analyses between the previously mentioned organoid and cell models using the DESeq2 package in R followed by principal component analysis (PCA) to identify variation within and between these datasets^10^. These analyses showed that the transcriptional signature of COs was highly similar to TOs but was distinct from both DOs and AECs (**Figure S2A-S2C**). COs differentially expressed 1322 transcripts relative to TOs (log_2_ fold change ± 2.5, p_adj_<0.01), with 769 differentially upregulated and 553 differentially downregulated (**Figure S2D, S2E, Table S1**). COs differentially expressed 2963 transcripts relative to DOs (log_2_ fold change ± 2.5, p_adj_<0.01), with 1208 differentially upregulated and 1755 differentially downregulated (**Figure S2D, S2G, Table S2**), and 2,240 transcripts relative to AECs, with 935 differentially upregulated and 1,305 differentially downregulated (**Figure S2D, S2F, Table S3**). K-means clustering and gene ontology (GO) enrichment analysis of differentially expressed transcripts between COs, TOs, DOs, and AECs revealed significant enrichment of pathways largely shared between COs and TOs that were distinctly enriched relative to DOs and AECs (**Figure 2A**). Significantly enriched pathways in COs and TOs included cytokine production (Cluster A, p=5×10^-4^), reproduction and pregnancy (Cluster B, p=5×10^-8^ and p=1×10^-9^, respectively), and gonadotropin secretion (Cluster B, p=3×10^-6^) (**Figure 2B**). Pathways specifically enriched in DOs included epithelial development (Cluster D, p=2×10^-6^), cell differentiation (Cluster D, p=5×10^-7^), and tissue development (Cluster E, p=3×10^-8^) (**Figure 2B**). AECs exhibited enrichment in cell adhesion (Cluster C, p=9×10^-18^) and animal organ development (Cluster C, p=2×10^-12^) (**Figure 2B**). Like TOs, COs highly expressed transcripts specifically enriched in human placental trophoblasts, including *CGA*, *CGB3*, PSGs (*PSG1/3/5/6/7*), *GATA3*, *XAGE2*, *XAGE3*, *PAGE4*, *ERVW-1*, and *SIGLEC6* which were absent or expressed at very low levels in DOs and AECs (**Figure 2C-E**, **Figure S2F-H**). As expected, DOs expressed transcripts previously shown to be enriched in these organoids^6^, including *SOX17*, *MUC5B*, and *FOXA2*, which were absent or expressed at low levels in COs, TOs, and AECs (**Figure 2C, 2F**). AECs also selectively expressed transcripts including *KRT5*, *KRT24*, and *IRX1*, which were not expressed in any organoid types (**Figure 2C, 2G)**.

**Figure 2:**
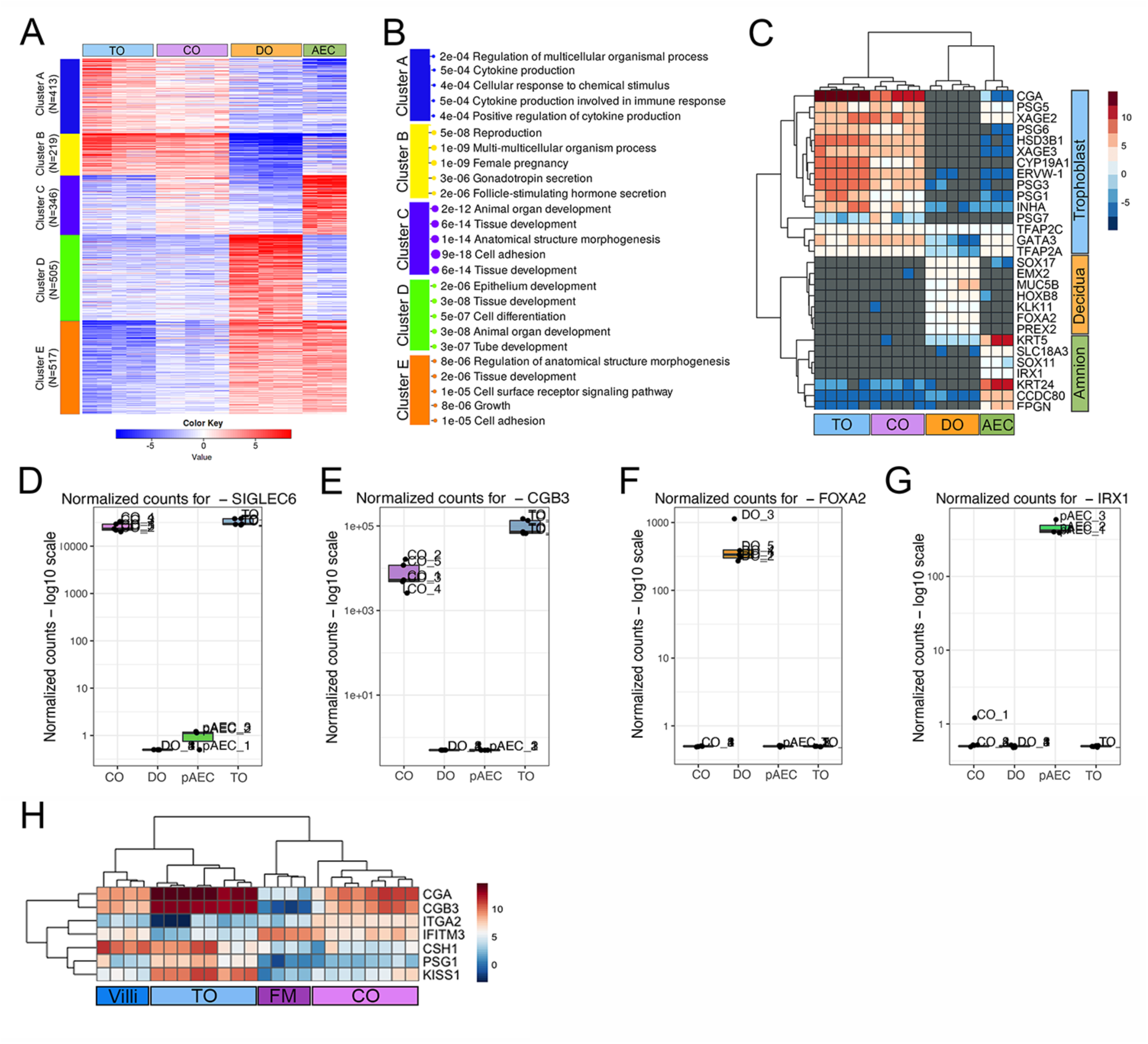
Transcriptional profiling of organoids and primary cells from distinct placental tissues by bulk RNA-seq. **(A),** K-means cluster analysis of GO Biological pathways enriched in villous-derived trophoblast organoids (TOs, blue at top), chorion-derived organoids (COs, purple at top), decidua organoids (DOs, orange at top) or primary amnion epithelial cells (AECs, green at top) based on differential expression analysis following RNA-seq. Five enriched clusters are shown (colored boxes at left). Key at bottom, with red indicating high values and blue indicating low values. TOs and COs were enriched for clusters A and B, COs and AECs were enriched for cluster C, DOs were enriched for cluster D, and DOs and AECs were enriched for cluster E. Numbers at left indicate genes associated with these clusters. **(B),** Select pathways associated with clusters A-E shown in A, with associated p-values shown at right. **(C),** Heatmap (based on log_2_ RPKM values) of transcripts expressed in TOs, COs, DOs, or AECs that are associated with markers of trophoblasts (top, blue), the decidua (middle, orange), or amnion (bottom, green). Key at right. Red indicates high levels of expression, blue indicates low levels of expression, and grey indicates no reads detected. Hierarchical clustering is shown on left and at top**. (D-G),** Normalized counts for the following transcripts in COs, DOs, AECs, and TOs: *SIGLEC6* (D), *CGB3* (E), *FOXA2* (F), and *IRX1* (G). Individual samples are shown as unique symbols. **(H),** Heatmap (based on log_2_ RPKM values) of the expression of the indicated transcripts in TOs (light blue) and COs (light purple) or mid-gestation chorionic villi (dark blue) and fetal membrane (FM, dark purple) samples.

We next compared the transcriptional profiles of COs and TOs to human placental tissue to determine if they recapitulated differential gene expression previously observed between chorionic villi and smooth chorion^2^. To do this, we performed bulk RNA-seq of chorionic villi and fetal membranes from four matched mid-gestation (16-21w) human placentas and compared the expression of select transcripts previously shown to be enriched in the smooth chorion^2^, including *ITGA2* and *IFITM3* as well as transcripts enriched in villi, such as *CSH1* and *CGA*. These analyses revealed that TOs clustered with chorionic villi, with both highly expressing *CSH1*, *CGA*, and other factors such as *CGB3* and *PSG1* (**Figure 2H**). Similarly, COs clustered with fetal membrane tissue, which expressed higher levels of both *ITGA2* and *IFITM3* (**Figure 2H**). These data collectively demonstrate that while COs express some transcripts conserved in its tissue of origin, COs broadly express canonical trophoblast markers similar to TOs.

### Single-cell RNA-Seq mapping of the cellular composition of villi- and smooth-chorion derived organoids

Villi-derived TOs recapitulate much of the trophoblast heterogeneity observed in human placental tissue^11^. To determine whether COs also recapitulate this heterogeneity, we performed single-cell RNA sequencing (scRNA-seq) on COs and TOs derived from three distinct placentas. Established lines of COs and TOs cultured/passaged for equivalent times were dissociated into single-cell suspensions and processed separately for sequencing and downstream analysis. Cellular suspensions were filtered, and cell viability was assessed before library preparation. Libraries were generated using a droplet-based method, and sequencing data were processed through a standard Seurat pipeline^12,13^. Following quality control steps—where cells with high mitochondrial gene expression or low gene counts were excluded—a total of 12,724 CO cells and 15,748 TO cells were retained for further analysis. These cells were integrated using Harmony to correct for batch effects across samples and sex regression was performed to control for sex-associated differences across samples^14^. Integration was followed by dimensionality reduction, including PCA and UMAP. Cluster analysis was performed using graph-based clustering, and cell type-specific marker expression was assessed for each cluster to assign putative cell identities. Cell-type annotation was done by comparing differentially expressed genes in each cluster to known placental cell markers. Clustering and cell-type specific enrichment analyses revealed seven major cell types in both TOs (**Figure 3A, 3B, S3A**) and COs (**Figure 3C, 3D, S3B**). These clusters included genes specifically enriched in markers of the STB (e.g., *CGB3*, *PSG3*, *INSL4*, *HOPX*, *HMOX-1*, *SDC1*), CTBs (e.g., *SLC27A2*, *PAGE4*, *PEG10*), proliferating CTBs (CTBp) (e.g., *MKI67*, *TOP2A*), and EVTs (e.g., *HLA-G*, *ITGA2*, *ITGB1*) (**Figure 3E, 3F**). In TOs, we identified four cluster of CTBp (CTBp-1-4), two clusters of CTBs (CTB-1 and CTB-2), and a single cluster mapping to the STB (**Figure 3A, 3E, 3G, S3C, Table S4**). Similarly, COs contained three CTBp clusters, two CTB clusters, and a single STB cluster (**Figure 3C, 3F, 3H, S3D, Table S5**). However, unlike TOs, COs also contained a cluster expressing genes associated with more mature EVTs (**Figure 3C, 3F, 3H, Table S5**). Collectively, these data define the cellular composition of chorion- and villi-derived organoids and show that these organoids both differentiate into multiple trophoblast lineages present in the human placenta.

**Figure 3.**
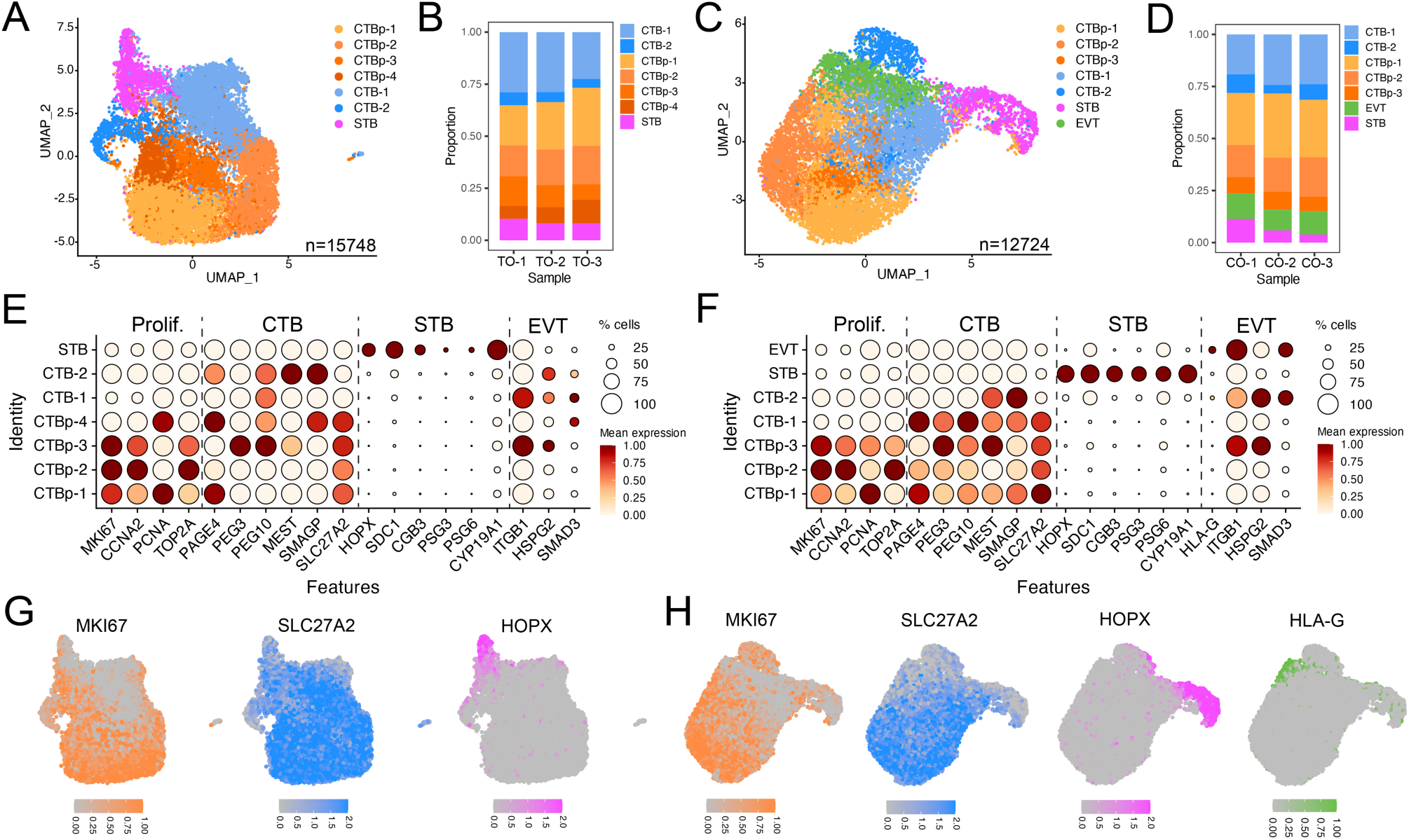
Single cell RNA-Seq of smooth chorion- and villi-derived organoids. **(A, C)**, UMAP of cell type clusters from three unique lines of villi-derived organoids (TOs, A, n=15748 cells) or chorion-derived trophoblast organoids (COs, C, n=12724 cells). **(B, D),** Comparison of the enrichment of distinct cell clusters in TOs (B) and COs (D) across individual samples. **(E, F),** Dot plots showing expression levels of indicated genes in each cluster of TOs (E) and COs (F). The indicated genes are established markers for proliferating cells (Prolif), cytotrophoblasts (CTBs), the syncytiotrophoblast (STB), and extravillous trophoblasts (EVTs). Scale is shown at right. **(G, H),** FeaturePlots of CTBp (MKI67), CTB (SLC27A2), STB (HOPX) and EVT (HLA-G) in cell clusters from TOs (G) or COs (H). Scale is shown at bottom.

### Comparative single-cell RNA-Seq analysis of villi- and smooth chorion-derived organoids reveals limited tissue-specific transcriptional enrichment

To determine whether there was differential enrichment of cell types and/or genes between COs and TOs, we performed comparative analyses using integrated datasets. This analysis revealed nine clusters, most of which were comprised of near-equivalent ratios of CO- and TO-derived cells (**Figure 4A, 4B, 4C, Figure S4A, S4B**). These clusters included three CTBp clusters, four CTB clusters, an STB cluster, and a single EVT cluster (**Figure 4A, 4B**)). Importantly, canonical markers of these trophoblast subpopulations were expressed at near equivalent ratios between COs and TOs (**Figure S4C, Table S6**). Differential expression analysis using DESeq2^10^ between CO- and TO-derived datasets identified the enrichment of 135 genes in CO-derived clusters and 56 genes in TO-derived clusters (**Figure 4D**). Genes enriched in COs were dispersed across all cluster/cell types, with differentially enriched genes (DEGs) most enriched in the EVT (18.66% of total) and CTB-3 (15.7% of total) clusters (**Figure 4E**). Several transcripts were enriched throughout all clusters, including Thrombospondin Type 1 Domain Containing 1 (*THSD1*), Nucleoredoxin Like 2 (*NXNL2*), and Family with Sequence Similarity 174 Member B (*FAM174B*) (**Figure 4F**). In contrast, other DEGs were enriched in a single cell cluster, including NLR Family Pyrin Domain Containing 7 (*NLRP7*) and *HLA-*G, both of which were enriched in the EVT cluster (**Figure 4F**). Additional DEGs included SRY-Box Transcription Factor 15 (*SOX15*) and the Cystatins *CTS1* and *CTS4*, all of which were enriched in both the EVT and CTB-3 clusters. (**Figure 4F, 4G**). The most highly enriched DEG in CTB-4 was *HLA-G* (**Figure 4G**). Taken together, these data further suggest that COs and TOs share many similarities, with limited transcriptional differences.

**Figure 4.**
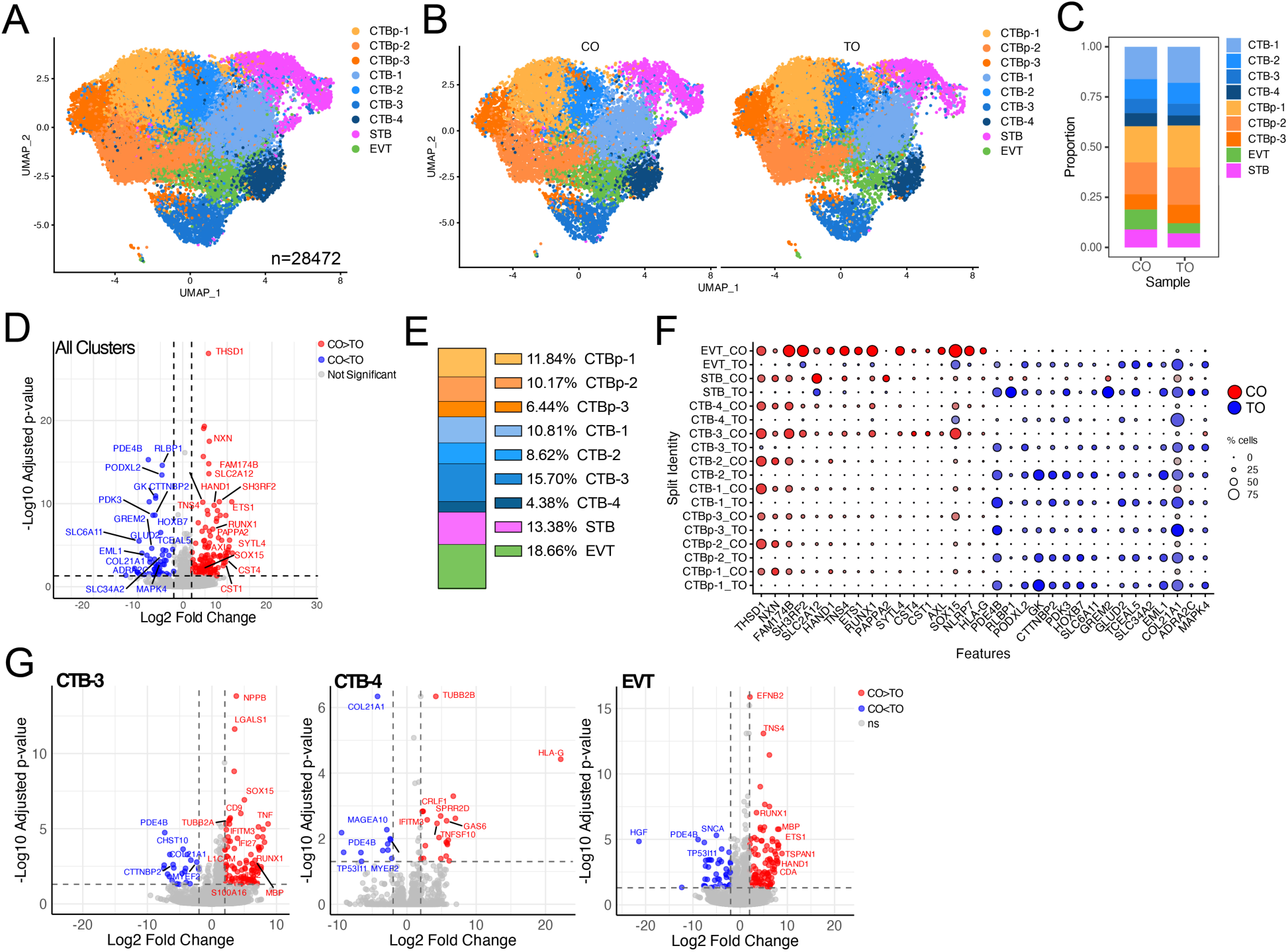
Comparative analysis of single-cell RNA sequencing of chorion- and villi-derived organoids. **(A)**, UMAP representation of cell type clusters from three distinct lines of villi-derived trophoblast organoid lines (TOs; A, n = 15,748 cells) and chorion-derived trophoblast organoid lines (COs; C, n = 12,724 cells), integrated into a single UMAP (n = 28,472 cells). **(B),** UMAPs of combined TOs and COs split by organoid type. **(C),** Comparison of the composition of distinct cell clusters from TOs and COs in the integrated dataset. **(D),** Volcano plot of the differentially expressed genes (DEGs) in all cell clusters between TOs and COs. Gene enriched in COs (CO>TO) are shown in red and those enriched in TOs (CO<TO) are shown in blue. Gene with no significant change are shown in grey. **(E),** Bar plot of the percent of DEGs in each cell cluster. **(F),** Dot plot showing the expression of DEGs enriched in COs (in red) or TOs (in blue) split by cell cluster and organoid type. Scale at right. **(G),** Volcano plot of the differentially expressed genes (DEGs) in the CTB-3 (left), CTB-4 (middle), and EVT (right) cell clusters between TOs and COs. Gene enriched in COs (CO>TO) are shown in red and those enriched in TOs (CO<TO) are shown in blue. Genes with no significant change are shown in grey.

### Comparative single-cell RNA-seq analysis of villi- and smooth chorion-derived tissue reveals extensive tissue-specific transcriptional enrichment

We identified several DEGs enriched in COs compared to TOs. To assess whether these DEGs were associated with their sites of origin, we first re-analyzed a publicly available scRNA-seq dataset from mid-gestation smooth chorion (SC) and chorionic villous (CV) tissues^2^. These datasets contained trophoblast populations including a single proliferating CTBp cluster, three CTB clusters, two EVT clusters, stromal cells, endothelial and epithelial cells, and several immune cell populations, which were present at similar ratios between SC and CV (**Figure S5A-S5C**). These clusters were enriched in canonical gene expression, which were expressed at near equivalent levels in SC and CV (**Figure S5D, S5E, Table S7**). To achieve better resolution of trophoblast populations, we subsetted the datasets to include only trophoblast cell types and performed re-clustering and UMAP analysis to more accurately capture the distinct transcriptional profiles of these cells. This resulted in seven distinct cell clusters, including one CTBp, three CTB populations, the STB, and three EVT populations (**Figure 5A**). Canonical cell type markers were expressed in both SC- and CV-derived clusters (**Figure S5H, Table S8**). Many of these cell populations were present at near-equivalent ratios in both SC and CV tissue, including CTBp, EVT-1, EVT-3, CTB-1, and CTB-3 (**Figure 5B, 5C, Figure S5F, S5G**). In contrast, three cell populations exhibited tissue-specific enrichment, which included the STB and EVT-2, which was more enriched in CV samples, and CTB-2, which was significantly enriched in the SC (**Figure 5B, 5C**).

**Figure 5.**
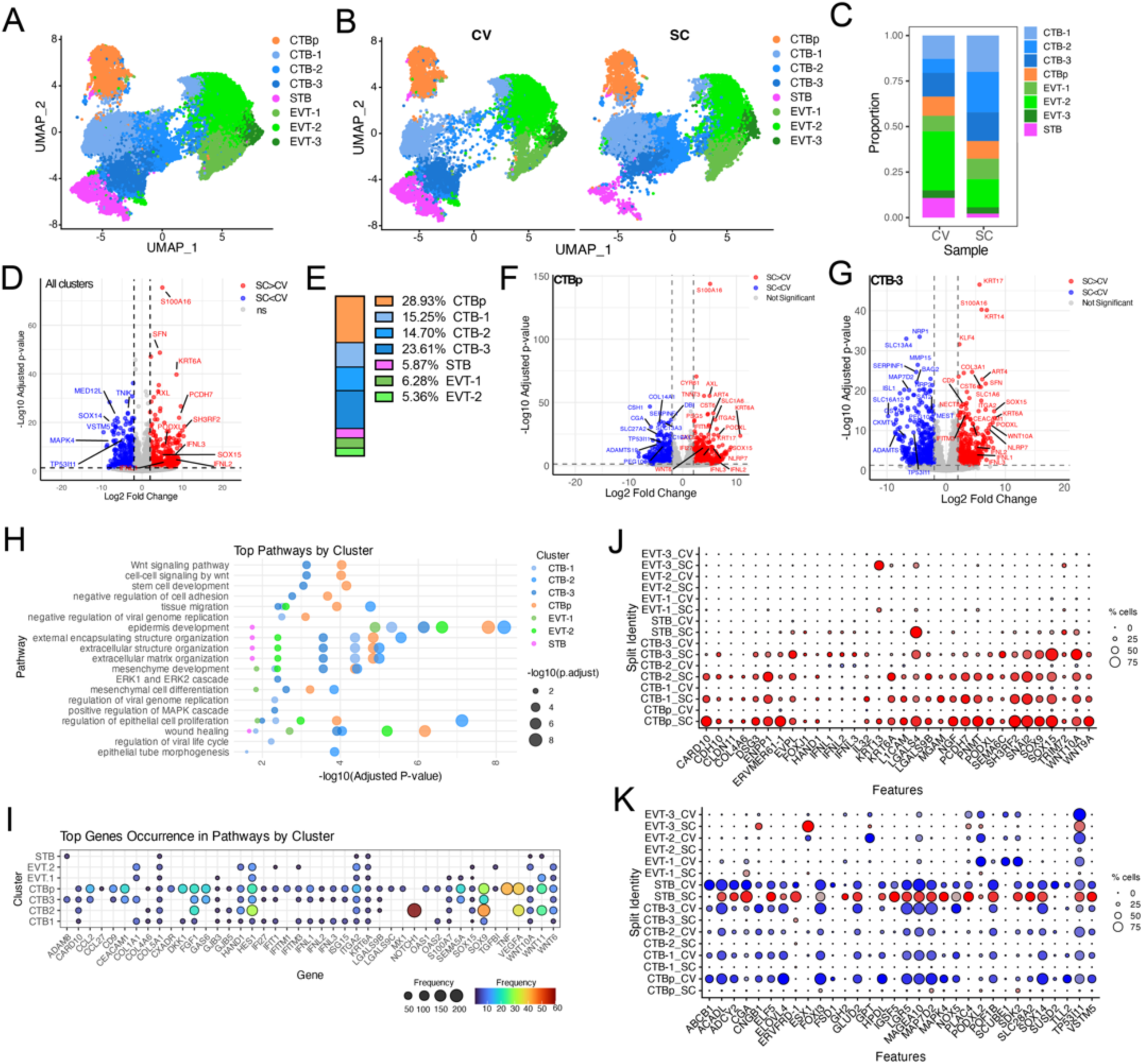
Comparative analysis of single-cell RNA sequencing of smooth chorion- and chorionic villi-derived tissue. **(A),** UMAP representation of trophoblast cell type clusters from both smooth chorion (SC) and chorionic villous (CV) tissue. **(B),** UMAP of combined CV and SC split by tissue type. **(C),** Comparison of the enrichment of distinct cell clusters from CV and SC in the integrated dataset. **(D),** Volcano plot of the differentially expressed genes (DEGs) in all cell clusters between CV and SC. Gene enriched in SC (SC>CV) are shown in red and those enriched in CV (SC<CV) are shown in blue. Gene with no significant change are shown in grey. **(E),** Bar plot of the percent of DEGs in each cell cluster. **(F, G),** Volcano plots of the DEGs in the CTBp (F) and CTB-3 (right), cell clusters between SC and CV. Gene enriched in SC (SC>CV) are shown in red and those enriched in CV (SC<CV) are shown in blue. Genes with no significant change are shown in grey. **(H),** DotPlot of GO pathway enrichment analysis indicating the pathways enriched in SC over CV arranged by cell cluster association. **(I),** Top genes associated with pathways shown in (H) and their expression in specific cell clusters. Key and scale at bottom. **(J, K),** Dot plot showing the expression of DEGs enriched in SC (in red) (J) or CV (in blue) (K) split by cell cluster and tissue type. Scale at right.

Next, we performed differential expression analysis using DESeq2, which identified 518 DEGs enriched in SC and 378 DEGs enriched in CV (**Figure 5D**). The majority of SC enriched DEGs were associated with the CTBp (28.93% of total DEGs) and CTB-3 (23.61% of total DEGs) populations (**Figure 5E**). This is consistent with a prior report indicating that there are differences in the progenitor cell population between CV and SC^2^. In contrast, all EVT populations and the STB contained ∼5% of total DEGs per cluster (**Figure 5E**). There were diverse transcripts enriched in the CTBp and CTB-3 clusters, which included several shared transcripts including all three-type III interferon (IFN) genes (*IFNL1-3*), *NLRP7*, *SOX15*, the cytokeratin 6A (*KRT6A*), and the S100 family member *S100A16* (**Figure 5G**). GO pathway analysis of all DEGs enriched in the SC showed enrichment of Wnt signaling and stem cell development in the CTBp and CTB-3 clusters, tissue migration in all CTB clusters, and regulation of viral genome replication and viral life cycle in select CTB clusters (**Figure 5H**). The most enriched genes associated with these pathways included IFNL genes and associated IFN stimulated genes (e.g., *IFITM1* and *IFITM3)*, *KRT6A*, the integrin *ITGA2*, *NOTCH1*, *SOX9*, *SOX15*, and several Wnt family members (*WNT10A*, *WNT11*, *WNT6*) (**Figure 5I**), which is consistent with previous reports^2^. The topmost enriched DEGs in the SC were heavily associated with CTB clusters (**Figure 5J**) whereas CV enriched transcripts in the STB were also present in the SC (**Figure 5K**). Collectively, these analyses highlight the significant differences in the transcriptional profiles of trophoblasts localized to distinct regions of the human placenta.

### Integrated Analysis Highlights Region-Specific Differences in Gene Expression in Organoids and Tissue from Smooth Chorion and Chorionic Villi

To further determine whether organoids recapitulated regional differences in trophoblast cell populations, we performed an integrated analysis of organoids derived from SC and CV alongside their respective *in vivo* counterparts. This approach allowed us to assess the similarities and differences between the organoids and native tissue. To do this, we integrated CO and TO datasets with trophoblasts from the SC and CV datasets described above and re-clustered the cells, generating a new UMAP to visualize associations more clearly. This dataset contained fifteen distinct cell populations, which included four CTBp clusters (CTBp1-4), five CTB clusters (CTB1-5), a pre-fusion STB cluster (STBpre), a single STB cluster, and four EVT clusters (EVT1-4) (**Figure 6A**), which expressed canonical cell type-specific markers (**Figure S6A, Table S9**). Many of these markers were expressed similarly between organoid and tissues, particularly in CTB and STB populations (**Figure S6B**). In contrast, EVT-associated markers including *HLA-G*, *SMAD3*, and *MMP-2* were expressed in organoids at significantly lower levels than in tissues (**Figure S6B**), which is expected given that trophoblast organoids lack significant spontaneous EVT differentiation in the absence of external factors^5,6^. Consistent with this, organoid models lacked significant enrichment in all EVT clusters despite similar levels of other cell types, including CTBp clusters and CTB clusters including CTB-1 and CTB-3, STBpre and the STB (**Figure 6B, 6C, Figure S6C**). PCA revealed that while COs and TOs were highly similar to each other, they formed distinct clusters separate from both CV and SC tissues (**Figure 6D**). Additionally, PCA showed that CV and SC tissues were distinct from each other, highlighting region-specific differences in gene expression and cellular composition (**Figure 6D**).

**Figure 6.**
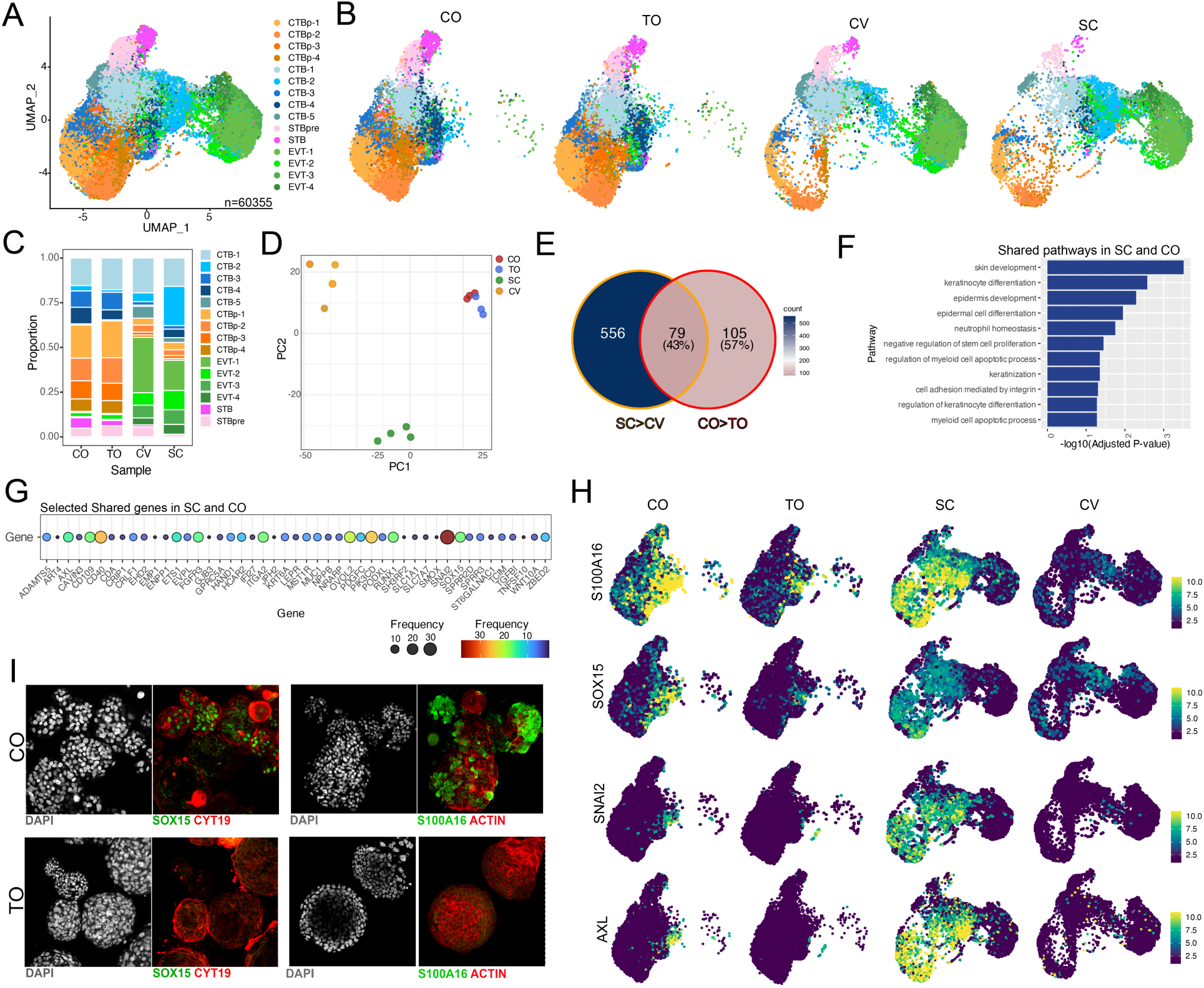
Single-cell profiling of smooth chorion and chorionic villous tissues and corresponding organoid types reveals commonly shared signatures with tissues of origin. **(A),** UMAP representation of cell type clusters from chorion organoids (CO), trophoblast organoids (TO), smooth chorion (SC) and chorionic villous (CV) tissue. **(B),** UMAP of combined UMAP in (A) split by sample type. **(C),** Comparison of the enrichment of distinct cell clusters from COs, TOs, CV, and SC in the integrated dataset. **(D),** Principal component analysis (PCA) of CO (red), TO (blue), CV (orange), and SC (green) datasets. **(E),** Venn diagram demonstrating the overlap of differentially expressed genes (DEGs) between SV and CV (SC>CV) datasets (orange outline) and CO and TO (CO>TO) datasets (red outline). Key is shown at right, with blue denoting the highest number of genes and red the lowest number. Shared gene (79 total) are shown in middle. **(F),** GO pathway enrichment analysis of shared DEGs between SC>CV and CO>TO datasets. **(G),** Top genes associated with pathways shown in (F) and their expression in specific cell clusters. Key and scale at bottom. **(H),** FeaturePlots of select genes differentially enriched in COs compared to TOs and SC compared to CV. Shown are S100A16 (top row), SOX15 (second row), SNAI2 (third row), and AXL (bottom row). Color scale at right. **(I),** Confocal microscopy for SOX15 (green, left panels) or S100A16 (green, right panels) and either cytokeratin-19 (red, left panels) or actin (red, right panels) in COs (top row) or TOs (bottom row). DAPI stained nuclei are shown in grey in all.

We next performed pseudobulk differential expression analysis between COs and TOs and SC and CV datasets to determine whether there were conserved gene signatures differentially expressed in a region-specific manner. We identified 184 differentially expressed genes in COs over TOs and 625 genes differentially enriched in SC over CV (**Figure 6E**). Of the genes enriched in COs, 43% were also differentially enriched in SC over CV (**Figure 6E**). Pathway analysis showed that these shared genes were associated with a number of pathways, including skin development, keratinocyte differentiation, epidermis development, negative regulation of stem cell proliferation, cell adhesion mediated by integrin, amongst others (**Figure 6F**). Genes associated with these pathways and shared in COs and SC datasets included *AXL*, *EVPL*, *ITGA2*, *OVOL2*, *RUNX1*, *SNAI2*, *SOX15*, and *TGFB1*, amongst others (**Figure 6G**). These genes were associated with specific clusters in COs and SC, particularly an enrichment in CTB-2 (**Figure 6H**). We confirmed the enrichment of selected transcripts, including SOX15 and S100A16, at the protein level through immunofluorescence microscopy, demonstrating their overexpression in COs compared to TOs (**Figure 6I**). Together, these data show that while COs and TOs exhibit significant similarity, there is a slight retention of tissue-associated transcriptional signatures, suggesting subtle distinctions in their molecular profiles.

### Microenvironmental Influence on Trophoblast Differentiation *in vivo* is Partially Recapitulated by Organoids

A previous study using scRNA-seq of matched CV and SC mid-gestation tissues suggested that the differentiation trajectories of trophoblasts at these sites are driven by distinct progenitor cell populations, each characterized by unique gene expression profiles and differentiation pathways^2^. To determine if these pathways were influenced by the *in vivo* tissue microenvironment, we performed comparative Slingshot trajectory analysis, a method that infers lineage trajectories by integrating pseudotime information, on the combined dataset of COs, TOs, CV, and SC. To do this, we generated a combined dataset in which all organoid and tissue types shared common cell clusters, which included CTBp-1-3, CTB1-3, STBpre, and STB clusters. (**Figure 7A, 7B, 7C**). Slingshot revealed three distinct trajectories, all originating from the CTBp-1 cluster. One of these trajectories extended from CTBp-1 to STB, passing sequentially through CTBp-3 and STBpre before culminating in the STB cluster (**Figure 7D**). A circular trajectory originated in CTBp-1 and cycled through all other CTBp clusters, including CTBp-2 and CTBp-3 (**Figure S7A).** Lastly, a trajectory originating in CTBp-1 passed through CTB-3 and CTB-1 before terminating in CTB-2 (**Figure 7E**). While all datasets contained the STB and CTBp trajectories, the CV sample did not contain a trajectory that terminated in CTB-2 (**Figure 7E**). Gene expression changes along these trajectories were then analyzed using fitGAM, which models gene expression dynamics across pseudotime to identify significant trends^15^. Gene expression along the CTBp-1 to STB trajectory was highly similar between organoids and tissues, with canonical STB-associated genes including *CSH1*, *CGB3*, *PSG3*, *HOPX*, *KISS1*, *CYP19A1*, and *PLAC4* and others increasing along the trajectory in all samples (**Figure 7F, 7H, Figure S7C**). Similarly, proliferation-associated genes such as *CENPF* underwent similar expression dynamics along the CTBp circular trajectory (**Figure S7B**). In contrast, gene expression along the CTBp-1 to CTB-2 trajectory was highly unique in the SC dataset and included the upregulation of distinct genes, some of which have been previously associated with schCTBs^2^, including *KRT6A*, *KRT14*, *KRT17*, *LAMA3*, and *COL5A1* and others (**Figure 7G, 7I, Figure S7D**). In some cases, these genes were also differentially expressed along the CTB-2 trajectory in COs, but not TOs. These included the schCTB marker *KRT6A* and transcripts associated with keratinocyte differentiation such as *KRT14* and *LAMA3* (**Figure 7G, Figure S7D**). Other transcripts enriched in both CO and SC along the trajectory included *S100A16* and *SOX15* (**Figure S7D**). Additional transcripts along the CTB-2 trajectory were exclusive to the SC, including *COL5A1* (**Figure S7D**).

**Figure 7.**
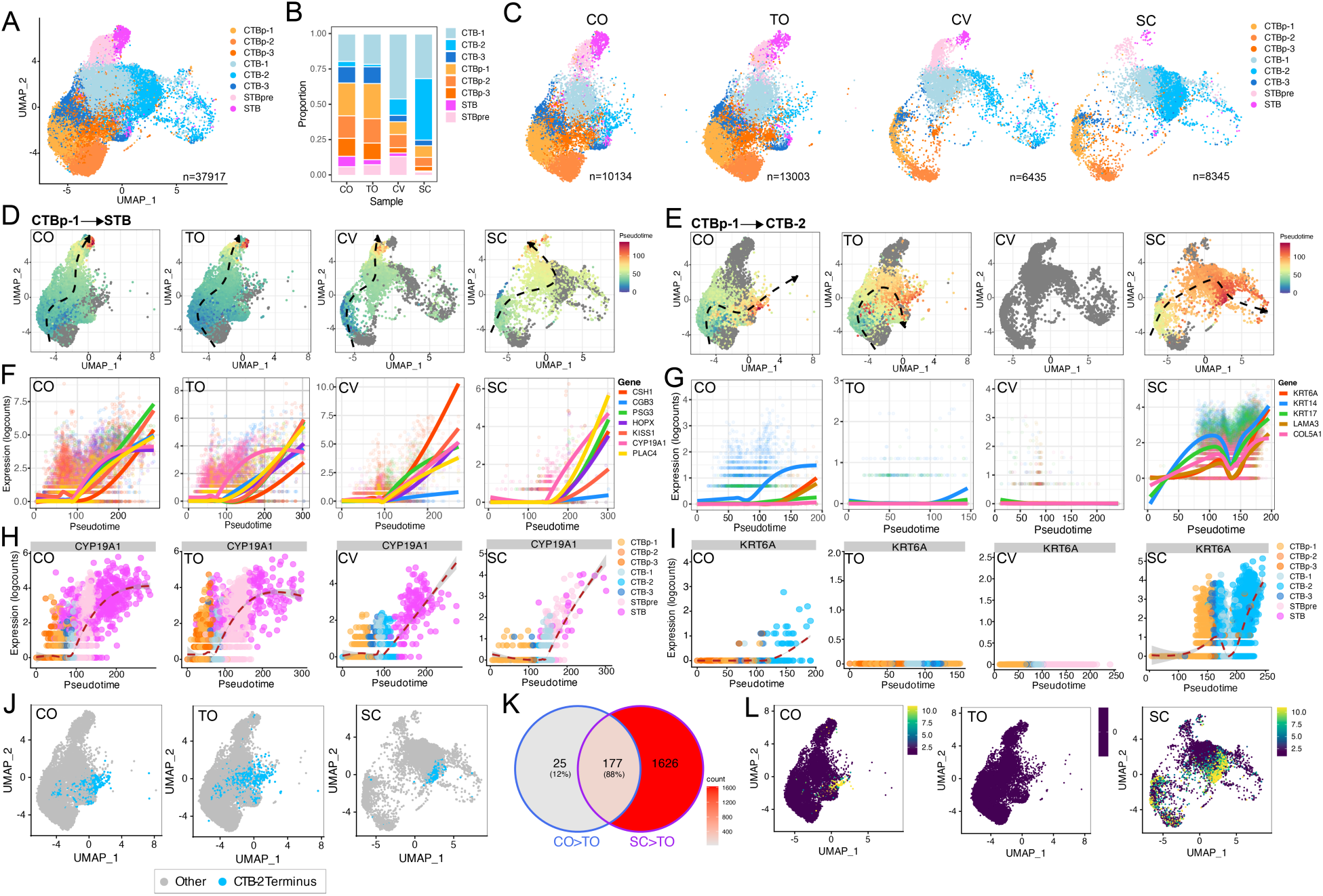
Comparative analysis of Trophoblast Differentiation across placental Tissues and respective Organoid types. **(A),** UMAP representation of trophoblast only cell clusters from chorion organoids (CO), trophoblast organoids (TO), smooth chorion (SC) and chorionic villous (CV) tissue. **(B),** Comparison of the enrichment of distinct cell clusters from COs, TOs, CV, and SC in the integrated dataset. **(C),** UMAP in (A) split by tissue type. **(D, E),** Trajectory analysis using Slingshot in COs, TOs, chorionic CV, or SC demonstrates differentiation pathways. The trajectory from a single cluster of proliferating cytotrophoblasts (CTBp-1) to the STB (D) or CTB-2 (E) is shown. The trajectory is color-coded from blue to red, indicating progression from the start to the end of the differentiation pathway. Black hatched lines trace the full trajectory, as calculated by Slingshot, with pseudotime inferred from lineage relationships. **(F, G),** Expression analysis of genes associated with the STB (F) or schCTBs (G) along the CTBp-1 to STB (F) or CTBp-1 to CTB-2 (G) pseudotimes using fitGAM. **(H, I).** Expression analysis of CYP19A1 (H) or KRT6A (I) along the CTBp-1 to STB trajectory (H) and CTBp-1 to CTB-2 trajectory (I) respectively using fitGAM. Red hatched lines represent the average gene expression along the trajectory, with pseudotime inferred using Slingshot. The color key on the right indicates the corresponding cluster identities. Expression patterns were calculated to highlight the changes along each differentiation path. **(J),** UMAP highlighting the terminal cells in the CTBp-1 to CTB-2 (in blue) trajectory. Cells in the final 10% of the Slingshot pseudotime are represented to indicate those reaching the end stage of these differentiation pathways. Note: CV does not contain a CTBp-1 to CTB-2 trajectory. Grey cells are not in the final 10% of the CTB-2 terminal trajectory. **(K),** Venn diagram denoting the overlap in genes shared in the terminal cells shown in (J) that were upregulated in COs over TOs (CO>TO) or SC over TO (SC>TO). Red scale indicates genes counts and percentages indicate total of those in the CO>TO gene list (scale at right). **(L),** FeaturePlots for KRT6A in COs (left panel), TOs (middle panel), or SC (right panel). Scale is shown at right, with yellow indicating higher gene expression.

To more comprehensively define differences in transcript expression in the terminal cells along the trajectories, we identified the terminal 10% of cells across all trajectories in the dataset (**Figure S7E**). These cells are expected to represent the final state of each cellular trajectory. We then conducted differential expression analysis between COs, TOs, and SC on these terminal states, with a particular focus on the CTB-2 lineage due to the SC-enriched characteristics of this trajectory (**Figure 7J**). This analysis revealed 202 differentially enriched genes in COs compared to TOs and 1803 genes in SCs compared to TOs (**Figure 7K**). Of the CO-enriched genes, 88% of them were conserved in the SC enriched gene list compared to TOs (**Figure 7K**). GO enrichment analysis revealed an enrichment of genes associated with skin development and keratinocyte differentiation (p<0.001) and included the schCTB marker *KRT6A* (**Figure 7L**). Collectively, these data suggest that while COs recapitulate certain aspects of schCTB differentiation, tissue-specific cues in the SC likely play a critical role in driving this differentiation.

## Discussion

Our work presented here establishes parallel trophoblast organoid models from placental chorionic villi and smooth chorion, providing a framework to explore the similarities and differences in trophoblast populations across these distinct sites. We demonstrate that COs and TOs differentiate to contain comparable trophoblast subpopulations and exhibit similar transcriptional profiles, despite originating from distinct placental sites. Using placental tissue datasets, we demonstrate that while chorion-derived organoids replicate certain aspects of schCTBs *in vivo*, they do not fully differentiate to recapitulate these cells, suggesting that the *in vivo* tissue microenvironment plays a crucial role in driving this differentiation. These studies provide a platform for advancing our understanding of trophoblast diversity and placental biology, offering valuable insights into the role of microenvironmental factors in trophoblast differentiation and the development of more accurate *in vitro* models for studying placental function and pathology.

By the end of the second month of pregnancy, extra-embryonic tissues segregate into two distinct regions that eventually form the villi-containing chorion frondosum and the uterine-facing region wherein villi degenerate to form the smooth chorion. Previous studies have shown that both schCTBs and villi-derived CTBs share expression of markers like HLA-G and epithelial-specific cytokeratins^1^. Similarly, whole-tissue explants from full-term and mid-gestation villous and fetal membrane tissues express and release similar cytokines and immune effectors^16,17^. However, an inherent limitation of these studies is their reliance on whole-tissue explants, which can obscure trophoblast-specific signaling or expression. Our work developed trophoblast organoid models to enable comparative studies of trophoblast populations derived from either chorionic villi or smooth chorion. Comparative transcriptional profiling of villi- and chorion-derived organoids reveals that the trophoblasts within these organoids share expression of many transcripts, including canonical trophoblast markers, hormones, and transcription factors exclusively expressed in trophoblasts. Differential expression analysis showed that while progenitor populations between COs and TOs are quite similar, EVT differentiation was notably different between the two, suggesting differences in the capacity of their progenitor cells to differentiate into EVTs. These findings imply that the local tissue environment may significantly shape the identity and function of these populations, contributing to their distinct characteristics *in vivo*. Notably, SOX15, a member of the SOX family, has been associated with trophoblast differentiation^18^, was differentially expressed in both COs and smooth chorion tissue. This differential expression of genes associated with stem cell differentiation underscores their potential role in regulating trophoblast progenitor cell states in response to specific environmental cues within the tissue, emphasizing the critical influence of the *in vivo* microenvironment in shaping progenitor cell behavior.

Chorionic villi and fetal membranes show distinct immune cell compositions and differing levels of interaction between trophoblasts and various fetal- and/or maternal-derived cells. For example, while chorionic villi contain fetal-derived Hofbauer cells, a specialized tissue-resident macrophage population responsible for modulating placental immune responses and engaging in tissue remodeling, these cells are absent in the fetal membrane. Chorionic villi also directly engage with various maternal-derived immune cell population, including maternal-derived macrophages/monocytes (PAMMs) and well as decidua NK cells (dNKs)^19,20^. In contrast, fetal membranes have a much sparser immune cell population, largely lacking resident macrophages and maternal-derived immune cells due to the physical separation from the maternal decidua. The primary immune cells found here are fetal in origin and include low numbers of macrophages and neutrophils, which are not involved in direct interactions with trophoblasts, as trophoblast invasion is minimal at this site. Consequently, fetal membranes maintain a more passive immune profile, emphasizing their role as a physical barrier without the active immune modulation seen in chorionic villi. Recent work has shown that specific cytokines and chemokines released from dNKs drive trophoblast differentiation in chorionic villi towards invasive EVTs^21^, highlighting the role that the *in vivo* environment plays in driving trophoblast differentiation. The organoid models described here thus provide a platform to study how the *in vivo* environment shapes trophoblast differentiation, enabling further exploration into the distinct immune profiles and differentiation pathways of trophoblasts across placental regions. By capturing key aspects of trophoblast behavior, these models offer insights into the dynamic interplay between trophoblasts and their microenvironment, particularly the influence of immune signals in guiding site-specific trophoblast differentiation.

Many of the shared genes and pathways between chorion-derived organoids and smooth chorion-derived tissue samples were associated with skin development and keratinocyte differentiation. This overlap underscores structural and functional similarities between the skin and fetal membranes, as both tissues are essential in forming and maintaining protective physical barriers. In both structures, specialized cell types and tightly regulated pathways work to create robust, selective barriers that protect against environmental pathogens while also maintaining essential permeability for molecular exchange. The shared pathways in skin development and keratinocyte differentiation highlight the evolutionary adaptation of fetal membranes to function much like the skin, reinforcing their role in safeguarding the fetus from external agents and physical stressors at the maternal-fetal interface. Notably, we observed differential expression of antiviral IFN-λ genes in the smooth chorion compared to chorionic villi, emphasizing its role as a protective barrier against ascending viral infections. These findings suggest that, akin to skin, the fetal membrane has evolved specialized mechanisms to defend against external threats, particularly at the maternal-fetal interface.

In summary, this study establishes parallel trophoblast organoid models from chorionic villi and smooth chorion, offering novel platforms to explore region-specific trophoblast differentiation and function. Our findings indicate that, although stem/progenitor-derived trophoblasts display a largely conserved gene expression profile across these regions, subtle transcriptional differences underscore the influence of the in vivo microenvironment on trophoblast identity and specialization. The partial conservation of region-specific gene expression patterns in organoids highlights the critical role of local environmental cues in driving the unique functions of trophoblast populations at distinct placental sites. These findings shed light on the complex relationship between trophoblasts and their surrounding environment, underscoring how tissue-specific cues are vital for preserving placental structure and promoting fetal health.

## Materials and Methods

### Human samples

Human tissue used in this study was obtained from Duke University after approval from the Duke University Institutional Review Board (IRB) and in accordance with the guidelines of Duke University human tissue procurement. Placental tissue used in this study for the generation of organoids was collected from third (38^th^-41^st^ weeks) trimester from C-sections. Placental tissue was excluded for diagnosis of complications including PPROM or maternal infection. Mid-gestation placental tissue (16-24 weeks) was obtained from the University of Pittsburgh Health Sciences Tissue Bank through an honest broker system after approval from the University of Pittsburgh IRB and in accordance with the University of Pittsburgh’s tissue procurement guidelines. Tissue was from healthy normal donors and tissue was excluded in cases of fetal anomalies or aneuploidy.

### Derivation and culture of organoids from human placental samples

Placental tissue was carefully dissected and separated into chorionic villi, decidua, chorion, and amnion separated. To remove the amnion layer, the amnion from intact fetal membranes was peeled from the chorion. To remove decidua from the chorion, isolated chorion tissue was carefully scraped with forceps. Stem/progenitor cell isolation from smooth chorion was similar to what has been described previously for full-term chorionic villous tissue ^6^. Briefly, dissected tissue was cut into small pieces (∼1-2mm^2^) and sequentially digested with 0.2% trypsin-250 (Alfa Aesar, J63993-09)/0.02% EDTA (Sigma-Aldrich, E9884-100G) and 1.0 mg/mL collagenase V (Sigma-Aldrich, C9263-100MG) on a 37°C stir plate in small bottles containing stir bars. Following the second digestion, tissue was disrupted manually by vigorous pipetting with a 5-10mL serological pipette (10-20 times). Nondigested tissue was removed by filtration over gauze and then pooled digests pelleted by centrifugation (600g, 6min) and then resuspended in ice-cold growth-factor-reduced Matrigel (Corning 356231). Matrigel “domes” (40 µl/well) were plated into 24-well tissue culture plates (Corning 3526), placed in a 37°C incubator to pre-polymerize for ∼3 min, turned upside down to ensure equal distribution of the isolated cells in domes for another 10 min, then carefully overlaid with 500 µL prewarmed media. For TOs and COs, cells were cultured in trophoblast organoid medium (TOM) consisting of Advanced DMEM/F12 (Life Technologies, 12634-010) supplemented with 1× B27 (Life Technologies, 17504-044), 1× N2 (Life Technologies, 17502-048), 10% FBS (vol/vol, Cytiva HyClone, SH30070.03), 2 mM GlutaMAX^TM^ supplement (Life Technologies, 35050-061), 100 µg/mL Primocin (InvivoGen, ant-pm-1), 1.25 mM N-Acetyl-L-cysteine (Sigma, A9165), 500 nM A83-01 (Tocris, 2939), 1.5 µM CHIR99021 (Tocris, 4423), 50 ng/mL recombinant human EGF (Gibco, PHG0314), 80 ng/mL recombinant human R-spondin 1 (R & D systems, 4645-RS-100), 100 ng/mL recombinant human FGF2 (Peprotech, 100-18C), 50 ng/mL recombinant human HGF (Peprotech, 100-39), 10mM nicotinamide (Sigma, N0636-100G), 5 µM Y-27632 (Sigma, Y0503-1MG), and 2.5 µM prostaglandin E2 (PGE2, R & D systems, 22-961-0). Cultures were maintained in a 37°C humidified incubator with 5% CO_2_. Medium was renewed every 2-3 days. To passage, COs digested in prewarmed TrypLE Express (Gibco, 12605-028) at 37°C for 10 min. Further mechanical dissociation was achieved by manual disruption. Dissociated CO fragments were collected and washed by centrifuge, then resuspended in fresh ice-cold Matrigel and replated as domes at the desired density for continuous culture. For freezing COs, overlaid media was aspirated, and organoid-Matrigel domes resuspended in CryoStor CS10 stem cell freezing medium (STEMCell Technologies, 07930) frozen at -80 ℃ and then transferred to liquid nitrogen for long-term storage. For thawing cryopreserved COs, organoids were thawed as quickly as possible, diluted with 5 times volume of basic TOM containing Advanced DMEM/F12, 2 mM GlutaMAX supplement, 10 mM HEPES (Gibco, 15630-106), 1 × Penicillin/Streptomycin (Lonza, 17-602E) and centrifuged to pellet. Afterwards, COs were resuspended in new ice-cold Matrigel and replated for recovery culture and passaged as described above. TOs and DOs were derived and maintained as described previously^6^. Conditioned media was was collected every 2-3 days. Media from at least ten timepoints for each established organoid line was collected as CO-CM or TO-CM. Non-conditioned medium (NCM) was tTOM (described above) that had not been exposed to organoids.

### Differentiation of COs and TOs into EVT-enriched organoids and Flow cytometry analysis

To generate EVT-enriched COs and TOs, we adopted EVT differentiation protocols developed previously^4,9^, including in TOs derived from full-term tissue^6^. Briefly, COs and TOs were passaged as described above and/or previously^22^ and plated directly into 8-well chamber slides (Millicell, EZslide, Millipore) pre-coated with ∼15 µl Matrigel or into Matrigel “domes”. Organoids were cultured in tTOM as described above for 3-4 days, then switched to EVT differentiation media 1 (EVT m1: advanced DMEM/F12 supplemented with 2 mM L-glutamine, 0.1 mM 2-mercaptoethenol, 0.5% (vol/vol) penicillin/streptomycin, 0.3% (vol/vol) BSA, 1% (vol/vol) ITS-X supplement, 100 ng/ml NRG1 (Cell Signaling, 5218SC), 7.5 µM A83-01, 2.5 µM Y27632 and 4% (vol/vol) Knockout Serum Replacement (Gibco 10828010), for 9 days. EVT m1 was renewed every 2 days for the initial 4 days then daily for the remaining culture period. After this period, organoids were switched to EVT m2, which is the same recipe as described above for EVT m1 but without NRG1, for a further 3-4 days, with daily media changes. Conditioned medium was collected as described below.

For flow cytometry analysis of EVT populations, EVT differentiation (EVTd) was performed in COs and TOs as descried above. After EVTd, two “domes” of organoids were collected and dissociated into single cells suspension by incubation with Stem Pro Accutase at 37 °C for 10 min then mechanical pipetting ∼400 times. Organoid single cell suspensions were incubated with PE-conjugated HLA-G (Abcam, ab24384, 1:100 dilution) or mouse isotype control (Thermo Fisher Scientific, 12-4714-42, 1:100 dilution) on ice for 30 min after filtration through a 100 μm strainer, Live/dead cell staining was performed using the LIVE/DEAD™ Cell Staining Kit (Thermo Fisher Scientific, L10119) on ice for 30 minutes, followed by blocking with human IgG (Sigma, I4506) at a concentration of 5 μg per 1 x 10⁶ cells on ice for 15 minutes. Cells were washed once with FACS buffer (1x PBS with 1% FBS [vol/vol]) before resuspension for analysis on the BD Accuri™ C6 Plus system according to the manufacturer’s instructions. Data analysis was conducted using FlowJo™ (v10.10.0), gating for live cells to assess cell surface HLA-G expression levels between COs_EVTd and TOs_EVTd samples.

### Derivation and culture of primary amnion epithelial cells from human fetal membrane

The amnion was physically separated from the chorion as described above. Amnion tissue was cut into small pieces and digested using the two-step protocol described above for chorion-derived organoids. Purity of each preparation was confirmed to be >90-95% based on immunostaining for the epithelial-specific cytokeratin cytokeraton-19 and actin followed by fluorescence microscopy. Cells were cultured in complete epithelial growth media (EpiCM, 4101, ScienCell Research Laboratories).

### Immunofluorescence microscopy

Organoids were fixed in 4% PFA for 15 min at room temperature, followed by 0.25% Triton X-100/PBS to permeabilize for 15 min at room temperature. Organoids were washed in 1x PBS and then incubated with the indicated primary antibodies in PBS at 4°C overnight. Organoids were pelleted by gravity, washed with PBS, and then incubated for 30 min at room temperature with Alexa Fluor-conjugated secondary antibodies (Invitrogen). Organoids were washed again with PBS and mounted in Vectashield (Vector Laboratories, H-1200) containing 4′,6-diamidino-2-phenylindole (DAPI) and were transferred onto microscope slides with large-orifice 200 µL-tips (Fisher Scientific, 02707134). The following antibodies and reagents were used: cytokeratin-19 (Abcam, ab9221, ab52625), HLA-G (Abcam, ab52454, ab283260), SOX15 (Invitrogen, 720155), S100A16 (Abcam, ab130419), Ki67 (550609, BD Biosciences, Alexa Fluor 594–conjugated phalloidin (Invitrogen, A12381), Alexa Fluor Plus 488 Goat anti-Mouse IgG secondary antibody (Invitrogen, A32723), Alexa Fluor 594 Goat anti-Mouse IgG secondary antibody (Invitrogen, A11032), Alexa Fluor 488 Goat anti-Rabbit IgG secondary antibody (Invitrogen, A11034), and Alexa Fluor 594 Goat anti-Rabbit IgG secondary antibody (Invitrogen, A11037). Images were captured using a Zeiss 880 Airyscan Fast Inverted or Olympus FV3000 confocal microscope and contrast-adjusted in Photoshop or Fiji. For whole Matrigel dome images, scans were performed using a 10x objective on an inverted IX83 Olympus microscopy with a motorized XY-stage (Prior) and tiled images automatically generated by CellSens (Olympus). Image analysis was performed using Imaris (version 9.2.1, Oxford Instruments).

### Immunohistochemistry

Tissue sections were deparaffinized with xylene and rehydrated with decreasing concentrations of ethanol (100%, 95%, 80%), then washed with ddH_2_O. Antigen retrieval was performed with slides submerged in 10 mM citrate buffer (pH 6.0) and heated in a steamer for 90°C for 20 min. Slides were cooled to room temperature and incubated with 6% H_2_O_2_ in methanol for 30 min. Following washing in 0.1% PBS-T (Phosphate-buffered saline, 0.1% Tween 20), slides were incubated in Avidin blocking solution for 15min, following by subsequent blocking in Biotin blocking solution for 15 min (Vector Laboratories, SP-2001). Following washing in 0.1% PBS-T, slides were then incubated with serum-free Protein Block (Abcam, ab156024) for 10min. Sections were incubated with primary antibody Ki67 (550609, BD Biosciences) diluted 1:250-500 in PBS-T overnight in a humidified chamber at 4°C. Next, slides were washed with PBS-T and incubated with secondary antibody (Biotinylated Mouse IgG, Vector Biolabs BA-9200) for 30min, washed, and then incubated with avidin/biotin-based peroxidase (Vector Laboratories, Vectastain Elite ABC HRP Kit, PK-6100) for an additional 30 min. Following washes in PBT-T, sections were incubated with DAB substrate (Vector Laboratories, SK-4100) for ∼5 min. Slides were washed with ddH_2_O and then counterstained with hematoxylin for 1 min, thoroughly rinsed with H_2_O, and incubated in 0.1% sodium bicarbonate in H_2_O for 5 mins. Slides were then dehydrated with increasing concentrations of ethanol, cleared with xylene and mounted with Vectamount Permanent Mounting Medium (Vector Laboratories, H-5000). Images were captured on an IX83 inverted microscope (Olympus) using a UC90 color CCD camera (Olympus).

### RNA extraction and Bulk RNA-seq

Total RNA was extracted with the Sigma GenElute total mammalian RNA miniprep kit following the manufacturer’s instruction and using the supplementary Sigma DNase digestion. RNA quality and concentration were determined using a Nanodrop ND-1000 Spectrophotometer. For bulk RNA-seq analysis, RNA was isolated from organoids as described above. Purified Total RNA was verified by Thermo scientific Nanodrop one. The libraries were prepared by the Duke Center for Genomic and Computational Biology (GCB) using the Tru-Seq stranded total RNA prep kit (Illumina). Sequencing was performed on the NovaSeq 6000 by using 150-bp paired-end sequencing. The reads were aligned to the human reference genome (GRCh38) using QIAGEN CLC Genomics (v20). Differential expression analysis was performed using the DESeq2 package in R ^10^, k-means pathway cluster enrichment for GO Biological processes and principal component analyses (PCA) was performed using iDEP ^23,24^. Heatmaps and hierarchical clustering was performed in R using the pheatmap package and were based on log_2_ RPKM (reads per kilobase million). Files associated with bulk RNA-seq studies have been deposited into Sequence Read Archive (PRJNA891648). Principal component analyses were performed using pcaExplorer in R^25^. Volcano plots were generated using the EnhancedVolcano package in R or in Graphpad Prism version 9.

### Dissociation of organoids for single cell RNA sequencing

Organoids were collected by carefully scraping Matrigel domes and transferring them into a 15 mL conical tube, followed by centrifugation. After removing the supernatant, 1 mL of pre-warmed TrypLE Express (Invitrogen, 12605036) was added, and the tube was incubated at 37°C for 12 minutes, with gentle swirling every 2-3 minutes. Dissociated organoids were pelleted at 1250 rpm for 3 min and re-suspended in 200 µL DMEM containing 10% FBS. Resuspended organoids were subjected to vigorous manual disruption using a single channel p200 pipette (Ranin, 17008652) for 3 min followed by the addition of 800 µL of DMEM containing 10% FBS. The dissociated suspension was then passed over a 40 µm filter cell strainer (Corning, 352098). Flow through was then centrifuged at 1250 rpm for 5min and the pellet resuspended in 250 µL of 1x PBS for a final volume of ∼300 µL and cell counts of ∼1 x 10^6^ cells/mL. Cell viability was determined using trypan blue and followed by automated cell counting (Biorad TC20 automated cell counter), with all samples containing ∼80% minimum viability. equencing was performed on three COs and TOs lines from unique placental tissues.

### Single cell RNA sequencing and analysis

Libraries were prepared from 10k total cells using the methods described in the 10x Genomics Single Cell 3’ Reagent Kit Protocol (v2 chemistry, Manual Part #CG00052). Sequencing runs were performed on an Illumina NovaSeq 6000 (Illumina, San Diego) using an S2 flow cell (Illumina, San Diego), which allowed for ∼61k reads/cell. Post-processing, quality control, and read alignment to the hg38 human reference genome were performed using 10x CellRanger package (v6.1.2, 10x Genomics). Gene expression matrices generated by the 10x CellRanger aggregate option were analyzed using Seurat (version 4.0) in R ^12,13,26,27^. Cells with at least 900 and no more than 12,000 unique expressed genes were included in downstream analysis, and cells with more than 25% mitochondrial reads were excluded from analysis. Donor sex was assigned using the biomaRt package in R^28^, based on the expression levels of sex-specific genes. Datasets from unique donors were normalized using the *SCTransform()* function^29^ in Seurat (version 4.4) in R. During normalization, variables such as the percentage of mitochondrial reads, number of features (nFeatures), number of counts (nCount), percentage of ribosomal reads, percentage of X-linked genes, and percentage of Y-linked genes were regressed out to mitigate their effects on downstream analyses. Dimensional reduction was initially performed using the *RunPCA()* function, with the first 30 principal components retained for downstream analyses. The number of components was selected based on the *ElbowPlots()* function, assessed across the first 50 dimensions. To correct for batch effects, Harmony batch correction was subsequently applied on these principal components^14^. To identify clusters, Louvain clustering (Seurat *FindClusters()* function) was performed, and optimal resolution was determined using the *clustree()* function^30^ on a range of resolutions between 0.2-1.0, with 0.4 used for downstream analyses. To identify clusters enriched in combined files of COs and TOs, datasets processed as described above were merged and Harmony applied to correct batch effects and align shared cell populations across the combined datasets. Clusters were defined as described above using a resolution of 0.4. Differential expression analysis between clusters was performed using the *FindAllMarkers()* function in Seurat with the Wilcoxon rank sum test. Parameters included a log₂ fold change threshold set to 0.25 and a minimum percentage of cells expressing the gene in either cluster set to 0.25. Genes with an FDR-adjusted p-value of < 0.05 were considered significant. Files associated with single cell RNA-seq studies have been deposited (GEO). For tissue analysis, publicly available datasets (GSE198373) described previously^2^ were downloaded and processed as described above, with cells with at least 800 and no more than 7500 unique expressed genes included in downstream analysis, and cells with more than 20% mitochondrial reads were excluded from analysis. Chorionic villi and smooth chorion datasets were normalized using *SCTransform* with the same parameters as described above, including regression of mitochondrial reads, ribosomal content, and sex-linked genes. Following normalization, datasets were integrated using Harmony to correct for batch effects and align shared cell populations across samples. Consistent parameters were used for dimensional reduction and clustering, with a resolution of 0.3 applied for tissue samples. A similar pipeline was implemented for integration of organoid and tissue datasets, with Harmony used for integration and clusters defined at a resolution of 0.4.

### Trajectory analysis

The *slingshot* package (version 2.6.0) in R was used to determine differentiation trajectories from clusters identified in Seurat setting the root of the trajectory in CTBp-1^31^. The raw counts and above-generated slingshot object were used to run *evaluateK()* with the total number of knots ranging from 3 to 9. The optimal number of knots was determined to be 5. The *fitGAM()* function using *tradeSeq* (1.5.10) was run with this resulting value and gene expression along lineages identified using the *associationTest()* function^15^. The *plotGeneExpression()* function was used to visualize raw count gene expression in individual cells across lineages from the *slingshot* object.

### Luminex assays

Luminex assays were performed using the following kits according to the manufacturer’s instructions: hCG Human ProcartaPlex Simplex Kit (Invitrogen, EPX010-12388-901) and Bio-Plex Pro Human MMP-2 Set (Bio-rad, 171BL029M). Plates were read on a Bio-Plex 200 (Bio-Rad) Luminex machines and analyzed using Bio-Plex (Biorad) software. Levels were averaged (in pg/mL unless otherwise stated) from at least three independent organoid lines.

### Statistics and reproducibility

All experiments reported in this study have been reproduced using independent samples (tissues and organoids) from multiple donors. All statistical analyses were performed using Prism software (GraphPad Software). Data are presented as mean ± SD, unless otherwise stated. Statistical significance was determined as described in the figure legends. Parametric tests were applied when data were distributed normally based on D’Agostino-Pearson analyses; otherwise, nonparametric tests were applied. For all statistical tests, p value <0.05 was considered statistically significant, with specific p values noted in the figure legends.

## Author contributions

LY designed and performed experiments, AC and CC designed and performed experiments and analyzed bulk and/or single cell sequencing datasets, and CC secured funding. All authors participated in manuscript review and writing.

## Acknowledgements

This work was supported by NIH AI145828 (C.B.C.). The authors are grateful to the patients who generously donated placental tissue for this research. We also thank Jennifer B. Gilner and Jillian Hurst for their efforts in performing and coordinating tissue collection, and Joshua Hatterschide for his assistance with scRNASeq analyses. We thank the Duke University School of Medicine for the use of the Sequencing and Genomic Technologies Shared Resource, which provided RNA-seq services. The authors also gratefully acknowledge the Duke Light Microscopy Core Facility for their technical support and assistance in this work.

