## Supplemental Figures for "Tissue Context Drives Regional Differences in Trophoblast Differentiation in Fetal Membranes and Chorionic Villi"

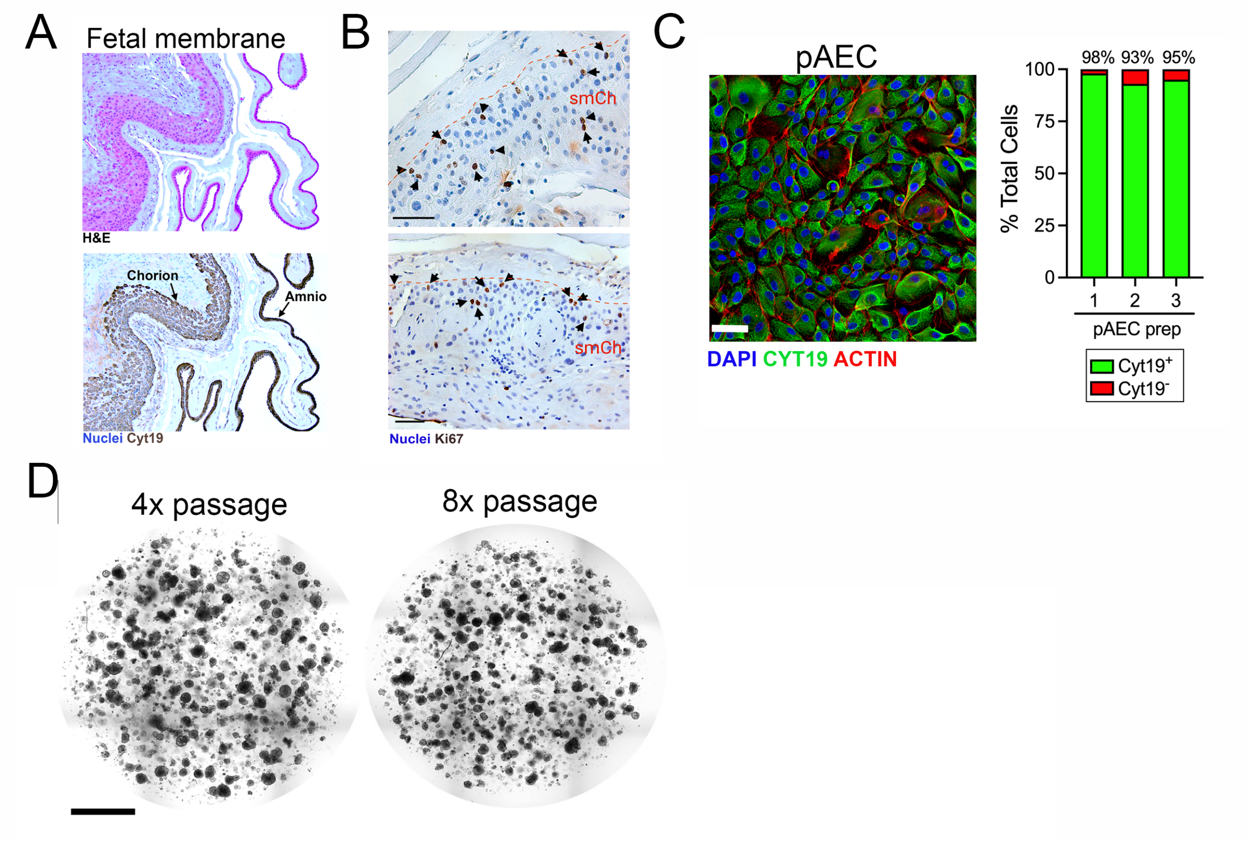


**Figure S1. Development of smooth chorion-derived organoids. (A),** H&E (top) or immunohistochemistry for cytokeratin-19 (in brown, bottom) of paraffin embedded whole fetal membrane sections isolated from full-term placentas used in this study. Arrows denote the amnion and chorion layers. **(B),** Immunohistochemistry for Ki67 (in brown) in the chorion layers (smCh) of two full-term fetal membrane tissues used in this study. Black arrows denote positive staining. Orange hatched line denotes the smooth chorion. Scale bar, 50μm. **(C),** Immunostaining for the epithelial cell-specific marker cytokeratin-19 (in green) and actin (in red) in primary amnion epithelial cells (AEC). DAPI-stained nuclei are in blue. Scale bar, 50μm. At right, quantification of three preparations of AECs showing the percentage of cells positive for cytokeratin-19 (in green). **(D),** Whole Matrigel dome brightfield images obtained by tile scanning of a representative line of chorion-derived organoids at either 4 (left) or 8 (right) passages post-isolation. Scale bar, 1mm.

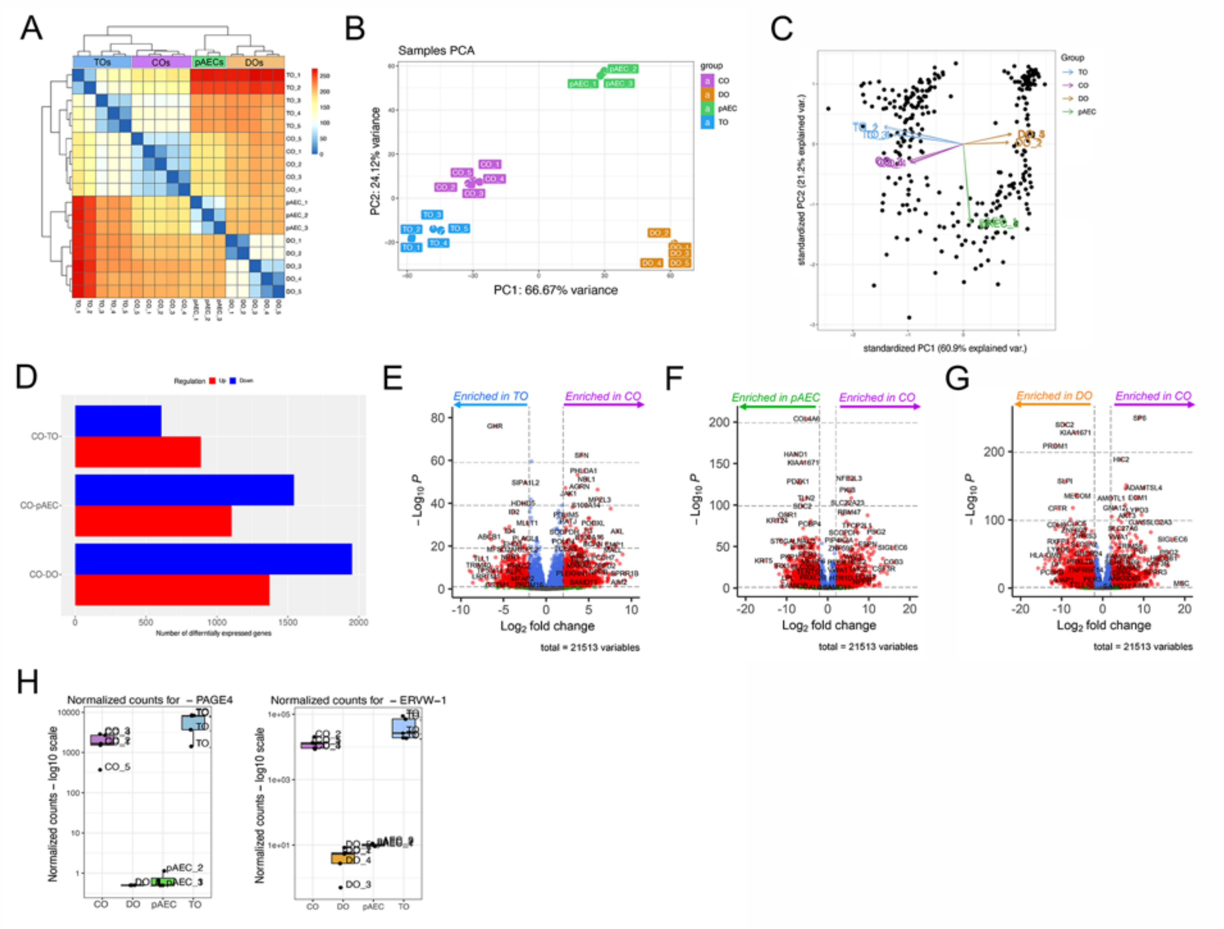

**Figure S2. Bulk RNA-seq analysis of placenta-derived organoids and primary cells. (A-C),** Principal component analysis of RNA-seq performed on villi-derived trophoblast organoids (TOs), decidua organoids (DOs), primary amnion epithelial cells (pAECs), and chorion-derived organoids (COs). In (A), heatmap of similarities between individual samples of TOs, COs, DOs, and pAECs. Key at right. In (B), PCA plotted on PC1 on x-axis and PC2 on y-axis of individual samples of TOs, COs, DOs, and pAECs. In (C), enrichment of transcripts in PC1 and PC2. **(D),** Differential expression of transcripts as determined by DeSeq2 in COs versus TOs (top), pAEC (middle), or DOs (bottom). Upregulated transcripts in red and downregulated in blue. **(E-G),** Volcano plots of the differentially expressed genes between COs and TOs (E), pAECs (F), or DOs (G). Purple arrow denotes genes expressed at higher levels in COs and blue (E), green (F), or orange (G), denote genes expressed at higher levels in TOs (E), pAECs (F), or DOs (G). **(H),** Normalized counts for *PAGE4* (left) and *ERVW-1* (right) in COs, DOs, pAECs, and TOs, Symbols represent individual replicates.

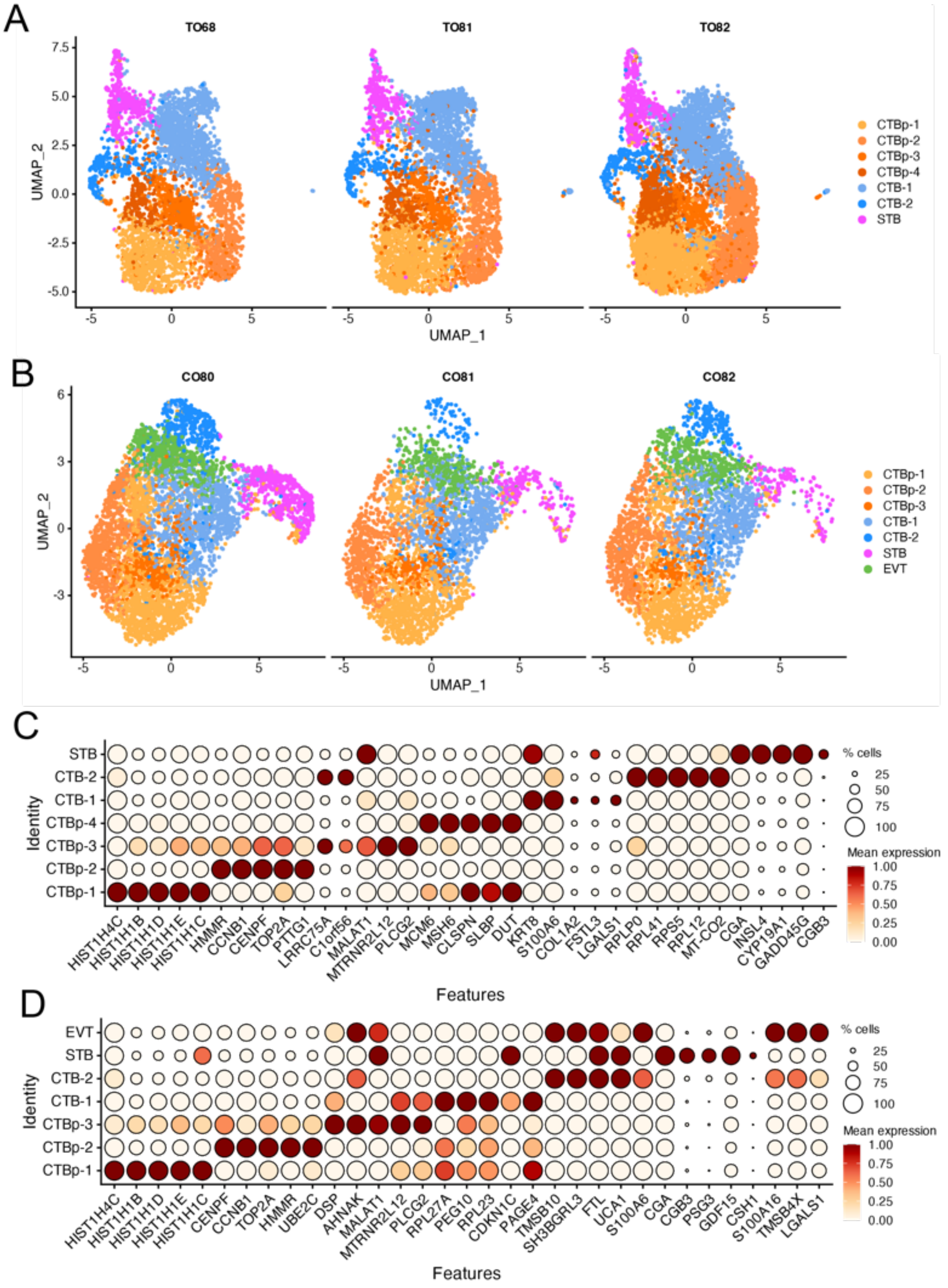


**Figure S3. Single-cell RNA-sequencing of COs and TOs. (A, B),** UMAPs of individual CO (A) or TO (B) lines separated by donor. **(C, D),** DotPlots of the top five markers genes per cluster in COs (C) or TOs (D). Scale is shown at right.

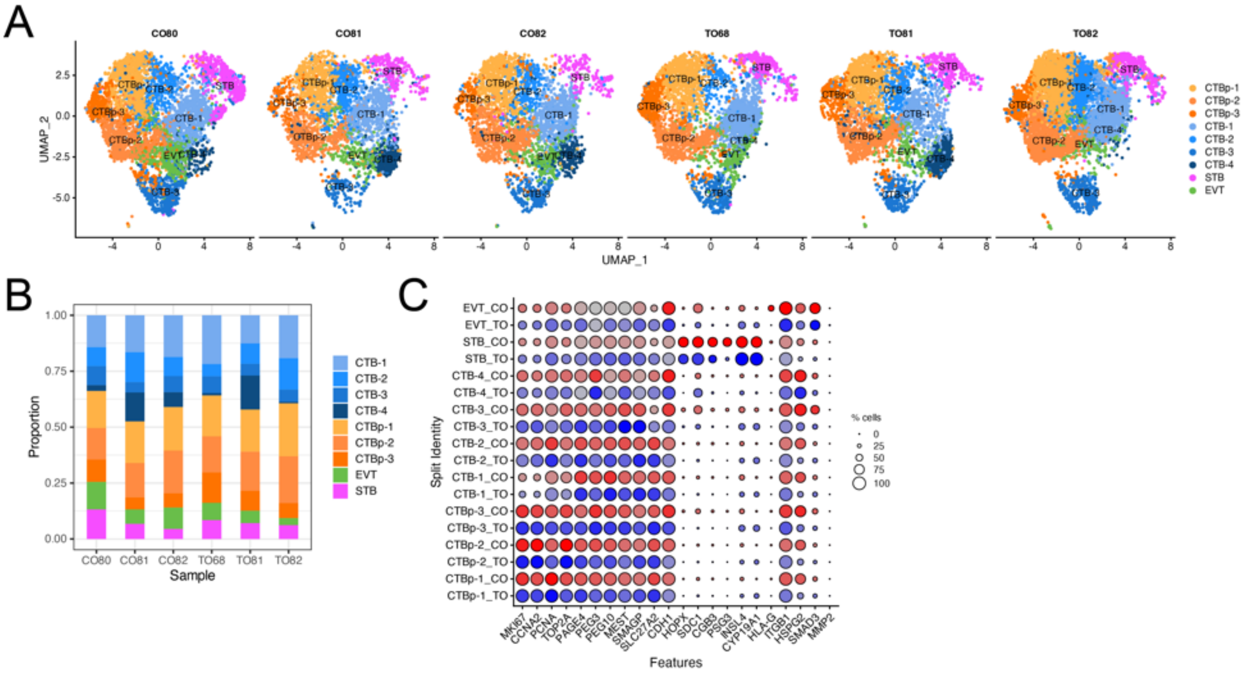


**Figure S4. Integrated single-cell RNA-sequencing of COs and TOs. (A),** UMAPs of individual CO or TO lines separated by donor. **(B),** Bar graph of cluster composition of individual CO and TO lines. **(C),** DotPlot of enriched cluster markers genes separated by organoid type (TO in blue and CO in red). Scale is shown at right.

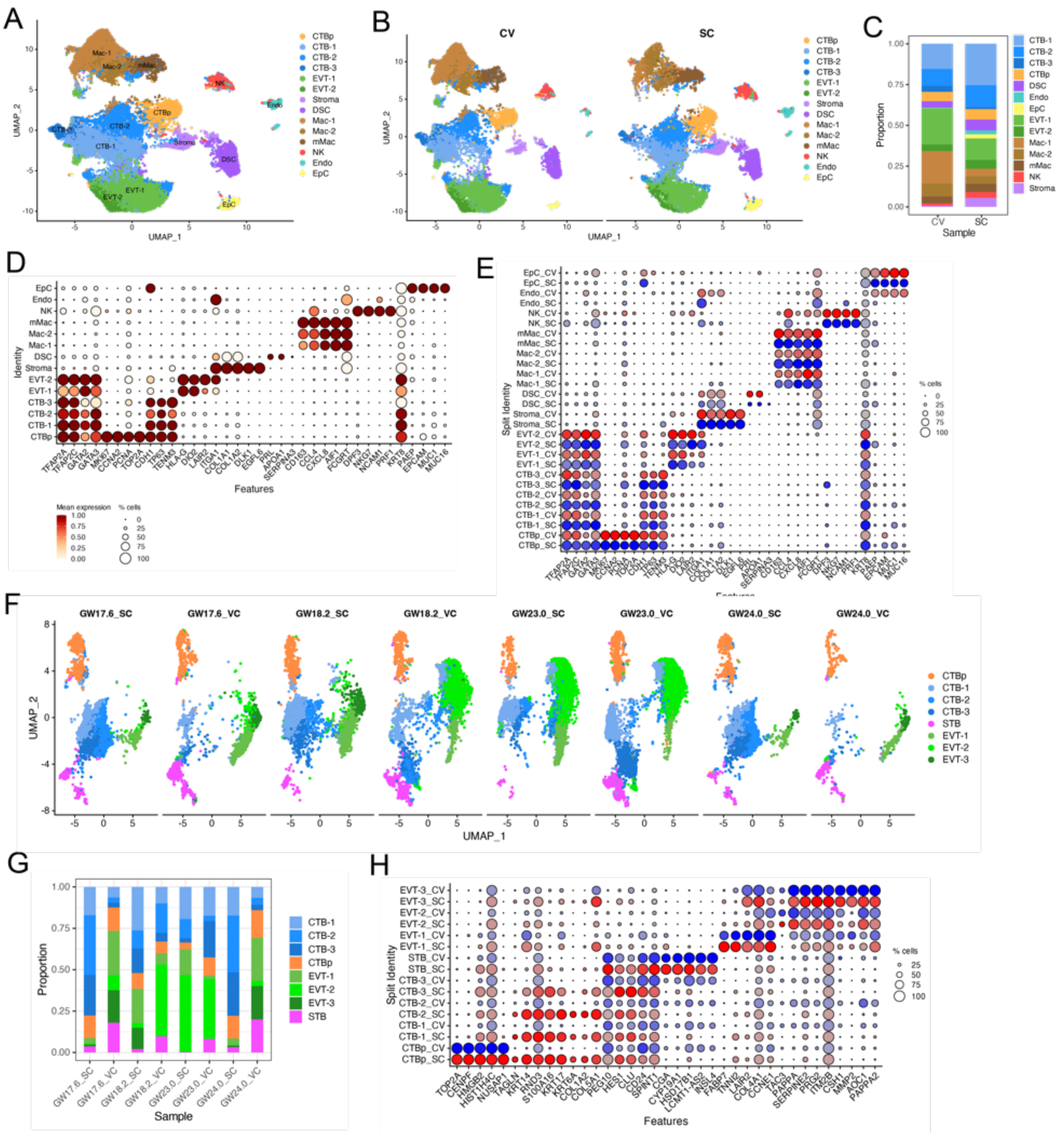

**Figure S5. Single-cell RNA-sequencing analysis of smooth chorion and chorionic villous tissue. (A, B),** UMAP of analyzed and integrated smooth chorion (SC) and chorionic villous (CV) datasets combined (A) or separated into tissue type (B). **(C),** Bar plot of cell clusters composition in SC or CV tissue. **(D, E),** DotPlots of enriched cluster marker genes in combined dataset (D) or separated by tissue type (SC in blue and CV in red). Scale is shown at bottom (D) or at right (E). **(F),** UMAPs of individual donor codes of trophoblast populations in SC or CV tissue. **(G),** Bar plot of the composition of trophoblast cell only clusters in individual donor codes of SC or CV tissues. **(H),** DotPlot of enriched cluster marker genes separated by tissue type (SC in red and CV in blue). Scale at right.

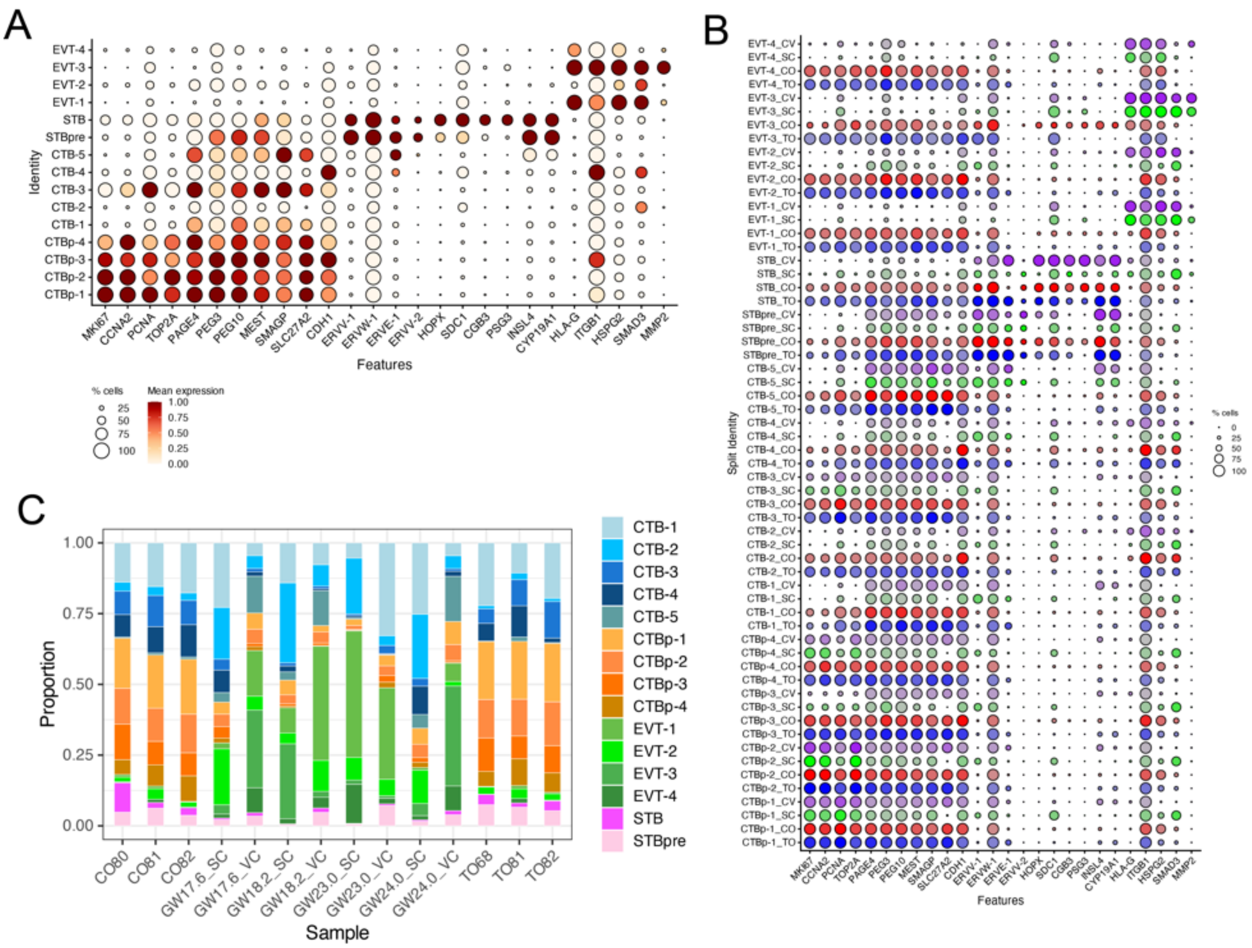


**Figure S6. Trophoblast populations in smooth chorion- and chorionic villous-derived organoids and corresponding tissues of origin. (A, B),** DotPlots of enriched canonical trophoblast linages marker genes in combined dataset (A) or separated by sample type (TO in blue, CO in red, SC in green, and CV in purple). Scale is shown at bottom (A) or at right (B). **(C),** Bar plot of trophoblast cell clusters composition in individual samples of COs, TOs, SC, or CV.

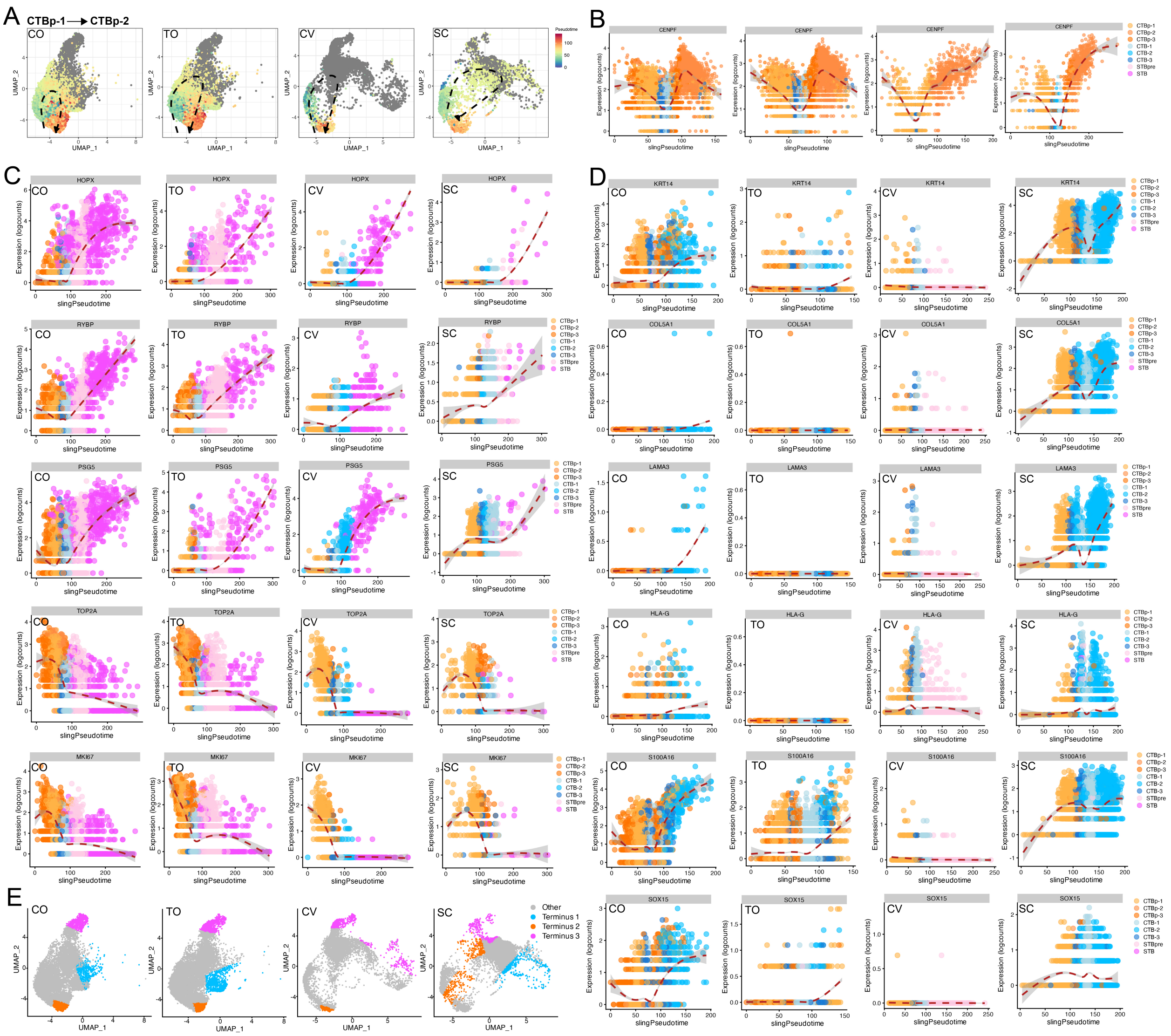


**Figure S7. Trajectory analysis in placental organoids and tissue. (A),** Trajectory analysis using Slingshot in chorionic organoids (COs), trophoblast organoids (TOs), chorionic villi (CV), or smooth chorion (SC) demonstrates differentiation pathways. The trajectory from a single cluster of proliferating cytotrophoblasts (CTBp-1) to CTBp-2 is shown. The trajectory is color-coded from blue to red, indicating progression from the start to the end of the differentiation pathway. Black hatched lines trace the full trajectory, as calculated by Slingshot, with pseudotime inferred from lineage relationships. **(B),** Expression analysis of the CENPF along the CTBp-1 to CTBp-2 trajectory **(C, D),** Expression analysis of the indicated genes along the CTBp-1 to STB trajectory (C) and CTBp-1 to CTB-2 trajectory (D) using fitGAM. Red hatched lines represent the average gene expression along the trajectory, with pseudotime inferred using Slingshot. The color key on the right indicates the corresponding cluster identities. Expression patterns were calculated to highlight the changes along each differentiation path. **(E),** UMAP highlighting the individual terminal cell populations from all three trajectories, including CTBp-1 to STB (in pink), CTBp-1 to CTB-2 trajectory (in blue), and CTBp-1 to CTBp-2 (in orange). Cells in the final 10% of the Slingshot pseudotime are represented to indicate those reaching the end stage of these differentiation pathways. Note: CV does not contain a CTBp-1 to CTB-2 trajectory. Grey cells are not in the terminal trajectories.
